# Pancancer survival analysis of cancer hallmark genes

**DOI:** 10.1101/2020.11.13.381442

**Authors:** Ádám Nagy, Gyöngyi Munkácsy, Balázs Győrffy

**Affiliations:** Department of Bioinformatics Semmelweis University Budapest, Hungary; TTK Momentum Cancer Biomarker Research Group, Budapest, Hungary

**Keywords:** gene expression, oncogenes, tumor suppressors, apoptosis, replicative immortality, induction of angiogenesis, metastasis, genomic instability, TCGA

## Abstract

Cancer hallmark genes are responsible for the most essential phenotypic characteristics of malignant transformation and progression. In this study, our aim was to estimate the prognostic effect of the established cancer hallmark genes in multiple distinct cancer types.

RNA-seq HTSeq counts and survival data from 26 different tumor types were acquired from the TCGA repository. DESeq was used for normalization. Correlations between gene expression and survival were computed using the Cox proportional hazards regression and by plotting Kaplan-Meier survival plots. The false discovery rate was calculated to correct for multiple hypothesis testing.

Signatures based on genes involved in genome instability and invasion reached significance in most individual cancer types. Thyroid and glioblastoma were independent of hallmark genes (61 and 54 genes significant, respectively), while renal clear cell cancer and low grade gliomas harbored the most prognostic changes (403 and 419 genes significant, respectively). The eight genes with the highest significance included BRCA1 (genome instability, HR=4.26, p<1E-16), RUNX1 (sustaining proliferative signaling, HR=2.96, p=3.1E-10) and SERPINE1 (inducing angiogenesis, HR=3.36, p=1.5E-12) in low grade glioma, CDK1 (cell death resistance, HR=5.67, p=2.1E-10) in kidney papillary carcinoma, E2F1 (tumor suppressor, HR=0.38, p=2.4E-05) and EREG (enabling replicative immortality, HR=3.23, p=2.1E-07) in cervical cancer, FBP1 (deregulation of cellular energetics, HR=0.45, p=2.8E-07) in kidney renal clear cell carcinoma and MYC (invasion and metastasis, HR=1.81, p=5.8E-05) in bladder cancer.

We observed unexpected heterogeneity and tissue specificity when correlating cancer hallmark genes and survival. These results will help to prioritize future targeted therapy development in different types of solid tumors.

## INTRODUCTION

Pancancer projects help to analyze the similarities and differences among different types of cancer by investigating genomic, epigenomic, transcriptomic and proteomic traits of the tumors. A leading effort in the pancancer genomic field is the PanCancer Atlas from the TCGA consortium ^1^, which focuses on the transcriptome, on the genomic interactions between somatic drivers and germline mutations, on the links to the methylome, on the proteome and on the tumor microenvironment and their implications for targeted and immune therapies ^2^.

During tumorigenesis, normal cells evolve to a neoplastic state in which they share common characteristics, including sustained proliferative signaling, loss of growth suppressors, apoptosis resistance, replicative immortality, angiogenesis induction, invasion and metastasis activation, genomic instability, inflammation, and energy metabolism reprogramming–the so-called “hallmarks of cancer” ^3 4^. A comprehensive database of genes associated with diverse cancer hallmarks was recently established, enabling the selection of hallmark-specific genes to be measured in transcriptome-level studies ^5^. Altogether, 671 cancer genes were grouped into eight main hallmark categories; notably, some of the genes were linked simultaneously to multiple hallmarks ^5^.

Analysis of gene expression contributed to the identification of molecular cancer subtypes capable of characterizing tumors and recognizing their biological characteristics, enabling the development of effectively targeted therapeutics. Single or multigene tests have been introduced to measure the deregulation of specific molecular pathways that can guide therapeutic decision-making by identifying genes that can serve as predictive or prognostic biomarkers. Breast cancer treatment is an outstanding example of a multigene decision tree-based treatment decision support protocol. The decision tree includes human epidermal growth factor receptor 2 (HER2), estrogen receptor (ER), and progesterone receptor (PgR). The overexpression or amplification of HER2 is present in approximately 25% of breast cancer cases ^6^. HER2-overexpressing tumors treated with anti-HER2 (trastuzumab and pertuzumab) therapy have improved disease-free and overall survival ^7^. ER-positive tumors are eligible for endocrine therapy ^8^. Increased disease-free and overall survival time was obtained by targeting ER with the antiestrogen tamoxifen in breast cancer ^9^. PgR positivity helps to improve the identification of ER-positive patients. ER, HER2, and PgR define three molecular subtypes of breast cancer, each with different treatment modalities. Those patients who are negative for all three markers are designated as triple-negative breast cancer; these patients have generally worse prognoses and conversely need a more aggressive systemic therapy.

Establishing prognostic multigene classification protocols can contribute to the understanding of tumor biology and to better prediction of cancer progression and cancer treatment strategies. One important issue is the selection of the proper method for the combination of the genes. First, genes can be utilized independently in a decision tree, where each node can be based on a single gene. Second, when multiple genes are combined, the most widespread approach is to compute their mean expression and to use this new value as a surrogate for the activity of the entire signature. A third option is to combine multiple genes after assigning a different weight to each of them. With breast cancer as an example, such combined signatures are utilized in FDA-approved multigene signature platforms, including the 76-gene signature, 21-gene signature and 70-gene signature platforms; all three of these can predict the prognosis of cancer under different conditions ^10 11 12^.

In this study, our goal was to rank established cancer hallmark genes according to their correlation to survival in a large cohort of distinct cancer types. We also aimed to correlate the relevance of each cancer hallmark in each of the available tumor types by assessing the prognostic power of signatures comprising hallmark genes.

## METHODS

### Database setup

All data processing steps and statistical analyses were performed in the R v3.5.2 statistical environment (http://www.r-project.org).

RNA sequencing (RNA-seq) data were utilized from the Cancer Genome Atlas (TCGA, https://cancergenome.nih.gov/). Only tumor types with more than 100 cancer specimens were included to ensure a robust sample number in each analysis.

The RNA-seq HTSeq count data generated by the Illumina HiSeq 2000 RNA Sequencing Version 2 platform were used in the expression analyses. The “DESeq” package based on the negative binomial distribution was used to normalize the raw count data ^13^. The Bioconductor “AnnotationDbi” package (http://bioconductor.org/packages/AnnotationDbi/) was applied to annotate Ensembl transcript IDs with gene symbols (n=25,228). A second scaling normalization was performed to set the mean expression of all genes in each patient sample to 1,000 to reduce batch effects.

For each sample, the preprocessed and annotated Mutation Annotation Format (MAF) data files that were generated by using MuTect2 for variant detection were used to compute the tumor mutation burden. The “maftools” package (http://bioconductor.org/packages/maftools/) was used for the aggregation and visualization of mutation data.

### Defining cancer hallmark signatures

Altogether, 671 cancer genes were grouped into eight hallmarks ^4^, based on gene assignment to hallmarks as described previously ^5^. The surrogate hallmark expression signature was calculated by computing the mean expression of all genes associated with the given hallmark in each tumor sample.

### Survival analysis and calculation of the strongest cutoff

Cox proportional hazards regression analysis was performed to examine the correlation between gene expression and overall survival (OS). The “survival” R package v2.38 (http://CRAN.R-project.org/package=survival/) was utilized to calculate log-rank *P* values, hazard ratios (HR) and 95% confidence intervals (CI). In addition, the survival differences were visualized by generating Kaplan-Meier survival plots.

To maximize the sensitivity of the analysis and to uncover any potential correlation to survival independent of a preset cutoff value (e.g., median), we computed each possible cutoff between the lower and upper quartiles of expression. Then, each of these cutoff values was used in a separate Cox regression analysis. The false discovery rate (FDR) was computed to correct for multiple hypothesis testing, and the result was only accepted as significant in the case of FDR<10%. The best performing cutoff with the lowest *P* value was used in the final analysis when drawing the Kaplan-Meier plot. The calculation of the best cutoff is demonstrated via the CDK1 gene in kidney papillary carcinoma and ovarian cancer in **Figure 1A** and **B**.

**Figure 1.**
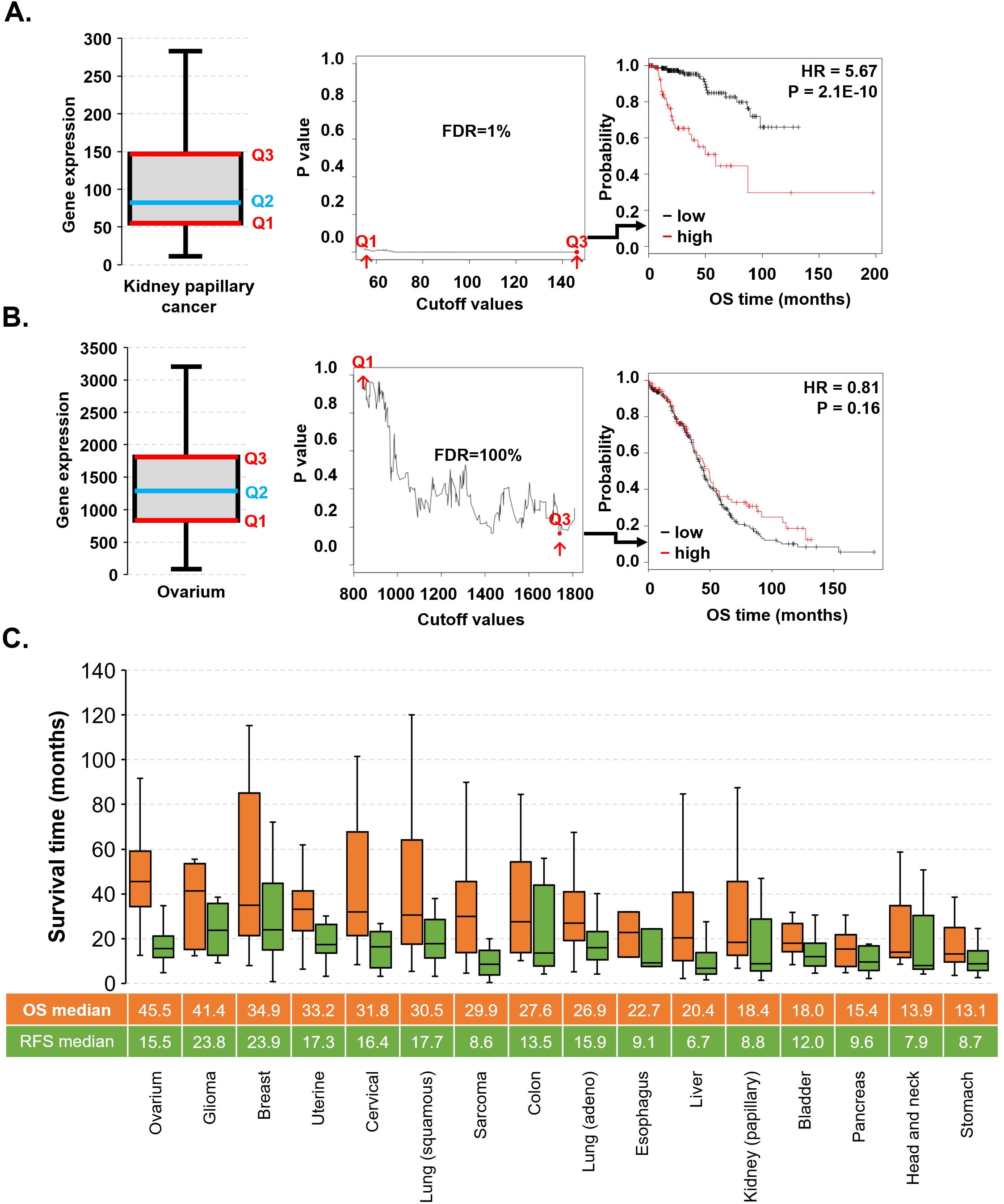
Overview of cutoff determination and survival distribution in the database. The determination of the best cutoff value in the survival analysis demonstrated with the CDK1 gene in kidney papillary carcinoma **(A)** and ovarian cancer **(B)**. Survival time characteristics of tumors with observed events **(C)**.

In addition, multivariate survival analysis was performed for the gene expression and clinical features to assess independence from known epidemiological and clinical variables, including race, sex, age, tumor stage and tumor grade.

### Data visualization

Hierarchical clustering was applied to group and to visualize the survival-associated cancer hallmark genes in different types of cancer using the Genesis software ^14^. The “forestplot” R package (https://CRAN.R-project.org/package=forestplot) was used to examine the association of cancer hallmark gene signatures with OS across different types of cancer. The “survplot” R package (http://www.cbs.dtu.dk/~eklund/survplot/) was used to generate the Kaplan-Meier plots.

## RESULTS

### Transcriptomic database

The complete dataset of RNA-seq samples with follow-up comprised 9,663 specimens from 26 distinct tumor types with breast cancer as the largest (n=1,090) and thymoma as the smallest set (n=118). Across the entire database, the median follow-up for overall survival (OS) was 24.3 months, and for relapse-free survival (RFS), it was 23.8 months. Most datasets contained both OS and RFS data, with the exception of AML, glioblastoma, melanoma and thymoma, which only had RFS data. Ovarian cancer patients had the highest median OS, while gastric and head and neck cancer patients had the shortest OS **(Figure 1C)**. In addition, glioma and liver cancer patients had the longest and the shortest median RFS at 23.8 and 6.7 months, respectively **(Figure 1C)**.

Clinico-pathological characteristics of patients, including stage, grade, sex and race, were available for 6,301, 4,126, 9,720 and 9,471 patients, respectively **(Table 1)**. According to the stage, head and neck cancer had the most patients in stage 4, and testicular cancer had the most patients in stage 0 or stage 1. The proportion of patients by tumor grade indicates that an unfavorable high grade was more common in bladder cancer, while a favorable low grade was restricted to head and neck cancer. Sex and ethnicity data of the patients showed that the number of males with cancer is higher than the number of females with cancer and that Caucasians give the majority in the TCGA database **(Table 1)**.

**Table 1.** Clinical characteristics of patients.

| Tumor type | TCGA code | Samples with RNA-seq data | Median survival – OS (months) | Events (n) | Median survival time in patients with an OS event | Median survival – RFS (months) | Events (n) | Median survival in patients with a relapse (months) | Sex (F/M) | Stage (S0/S1/S2/S3/S4) | Grade (Low/High) | Race (White/Asian/Black -African) |
| --- | --- | --- | --- | --- | --- | --- | --- | --- | --- | --- | --- | --- |
| AML | LAML | 151 | 10.13 | 97 | 7.13 | 0.00 | 0 | - | 68/83 | - | - | 135/1/13 |
| Bladder | BLCA | 405 | 17.87 | 179 | 13.60 | 0.00 | 31 | 15.40 | 106/299 | 0/2/130/138/133 | 21/381 | 321/44/23 |
| Breast | BRCA | 1090 | 28.10 | 151 | 42.40 | 21.35 | 84 | 25.77 | 1078/12 | 0/181/619/247/20 | - | 752/61/182 |
| Cervical | CESC | 304 | 21.23 | 71 | 20.23 | 12.75 | 26 | 16.10 | 304/0 | - | 153/119 | 209/20/30 |
| Colon | COAD | 454 | 22.30 | 102 | 13.47 | 0.00 | 23 | 16.87 | 214/240 | 0/75/176/128/64 | - | 212/11/59 |
| Esophagus | ESCA | 161 | 13.57 | 64 | 13.38 | 0.00 | 21 | 7.47 | 23/138 | 0/16/69/49/8 | 82/44 | 100/38/5 |
| Glioblastoma | GBM | 153 | 11.90 | 122 | 12.70 | 0.00 | 1 | 51.67 | 54/99 | - | - | 137/5/10 |
| Glioma | LGG | 510 | 22.12 | 125 | 27.13 | 0.00 | 20 | 19.93 | 228/282 | - | 248/261 | 470/8/21 |
| Head and neck | HNSC | 500 | 21.27 | 217 | 14.33 | 0.00 | 28 | 7.70 | 133/367 | 0/25/70/78/259 | 360/121 | 426/10/47 |
| Kidney (clear cell) | KIRC | 530 | 39.85 | 173 | 27.30 | 0.00 | 15 | 30.00 | 186/344 | 0/265/57/123/82 | 241/281 | 459/8/56 |
| Kidney (papillary) | KIRP | 288 | 25.58 | 44 | 21.37 | 13.22 | 28 | 15.72 | 76/212 | 0/172/21/51/15 | - | 205/6/60 |
| Liver | LIHC | 371 | 19.57 | 130 | 13.85 | 10.73 | 143 | 9.10 | 121/250 | 0/171/86/85/5 | 232/134 | 184/158/17 |
| Lung (adeno) | LUAD | 513 | 21.13 | 187 | 19.93 | 9.80 | 89 | 15.90 | 276/237 | 0/274/121/84/26 | - | 387/7/52 |
| Lung (squamous) | LUSC | 501 | 21.63 | 216 | 17.85 | 11.83 | 61 | 18.40 | 130/371 | 0/244/162/84/7 | - | 349/9/30 |
| Melanoma | SKCM | 468 | 34.45 | 215 | 35.67 | 0.00 | 0 | - | 179/289 | 7/76/140/170/23 | - | 445/12/1 |
| Ovary | OV | 374 | 34.03 | 230 | 36.55 | 0.00 | 126 | 17.67 | 374/0 | - | 43/321 | 324/11/25 |
| Pancreas | PAAD | 177 | 15.43 | 92 | 12.90 | 0.00 | 23 | 14.97 | 80/97 | 0/21/146/3/4 | 125/50 | 156/11/6 |
| Paraganglioma | PCPG | 178 | 25.08 | 6 | 15.08 | 20.42 | 4 | 27.65 | 101/77 | - | - | 147/6/20 |
| Prostate | PRAD | 495 | 30.80 | 10 | 36.73 | 20.53 | 30 | 25.30 | 0/495 | - | - | 147/2/7 |
| Rectum | READ | 165 | 20.33 | 25 | 20.33 | 0.00 | 6 | 28.68 | 75/90 | 0/30/51/51/24 | - | 80/1/6 |
| Sarcoma | SARC | 259 | 31.57 | 98 | 22.27 | 5.37 | 66 | 11.17 | 141/118 | - | - | 226/6/18 |
| Stomach | STAD | 375 | 14.23 | 147 | 11.60 | 6.60 | 37 | 10.50 | 134/241 | 0/53/111/150/38 | 147/219 | 238/74/11 |
| Testis | TGCT | 134 | 42.03 | 4 | 18.85 | 20.67 | 27 | 15.03 | 0/134 | 0/55/12/14/0 | - | 119/4/6 |
| Thymoma | THYM | 119 | 38.83 | 9 | 28.43 | 0.00 | 0 | - | 57/62 | - | - | 99/12/6 |
| Thyroid | THCA | 502 | 31.47 | 16 | 34.03 | 18.72 | 26 | 16.43 | 367/135 | 0/281/52/112/55 | - | 332/51/27 |
| Uterine | UCEC | 543 | 30.37 | 91 | 23.63 | 21.03 | 57 | 17.33 | 543/0 | - | 218/325 | 372/20/106 |
| Σ | - | 9,720 | 24.33 | 2,821 | 19.23 | 23.8 | 972 | 15.6 | 5,048/4,672 | 7/1,941/2,023/1,567/763 | 1870/2256 | 7,031/596/844 |

### Prognostic significance of hallmark-associated genes across 26 types of cancer

Cox regression analysis was performed using the RNA-seq expression of 671 cancer hallmark genes. The results of survival analysis across 26 types of cancer for each gene are listed in **Supplemental Table 1**. We computed the proportion of significant genes in each hallmark and in each tumor type **(Figure 2)**. Hierarchical clustering was performed to correlate different tumor types and cancer hallmark-associated genes. In this analysis, genes associated with invasion and metastasis activation, genome instability, sustained proliferative signaling and cellular energetics deregulation clustered into separate cohorts **(Figure 2)**. The top five tumors that contained the highest proportion of established cancer hallmark genes significantly associated with overall survival were kidney renal clear cell carcinoma, low grade glioma, melanoma, thymoma, and liver cancer.

**Figure 2.**
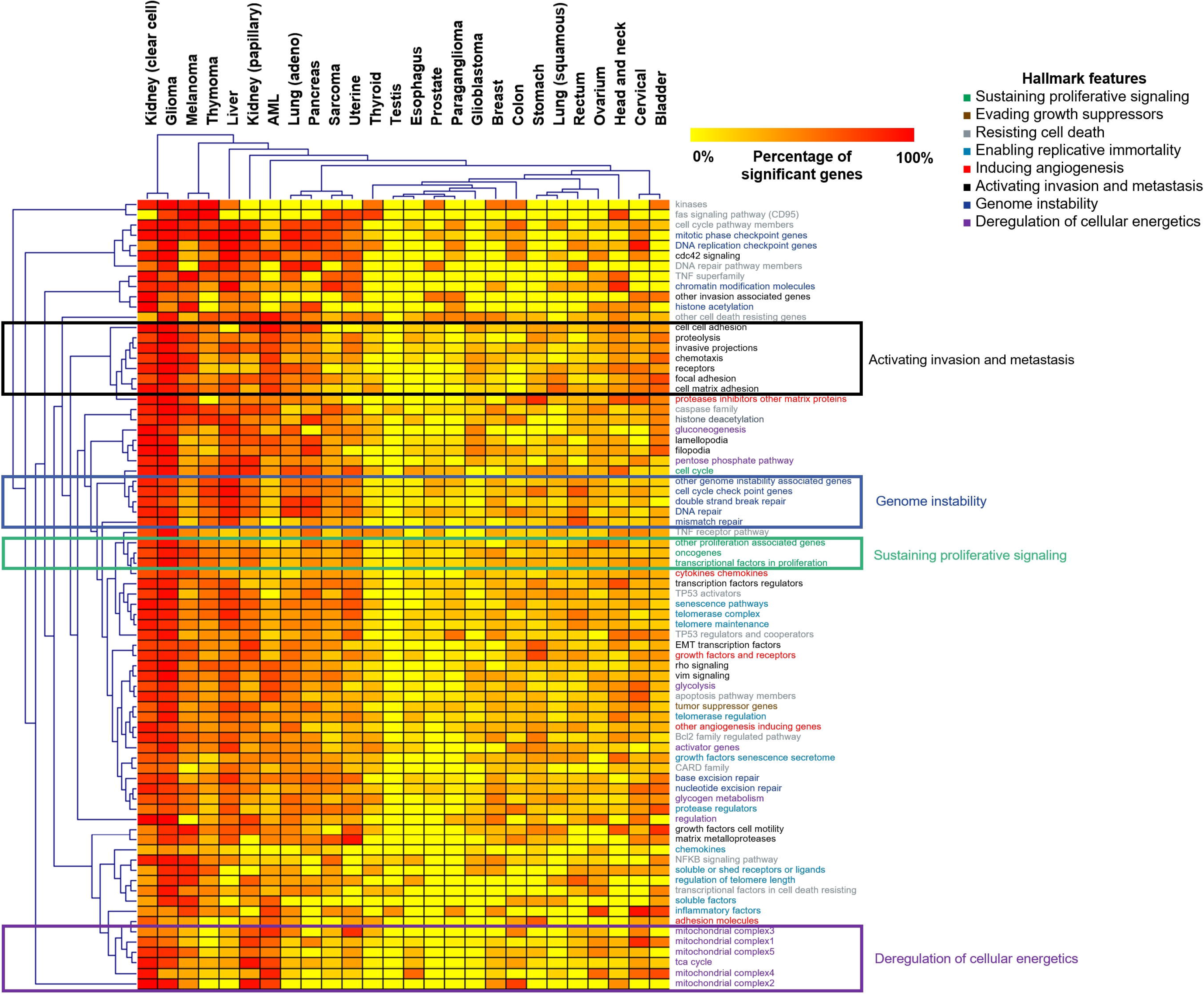
The prognostic power of cancer hallmark genes.

### Hallmark signatures and survival in different types of tumors

The expression signature of hallmark features was determined for each sample, and the prognostic effect of these signatures was investigated in different types of cancer. Significant *P* values (*P*<0.05) are illustrated as forest plots in **Figure 3A**.

**Figure 3.**
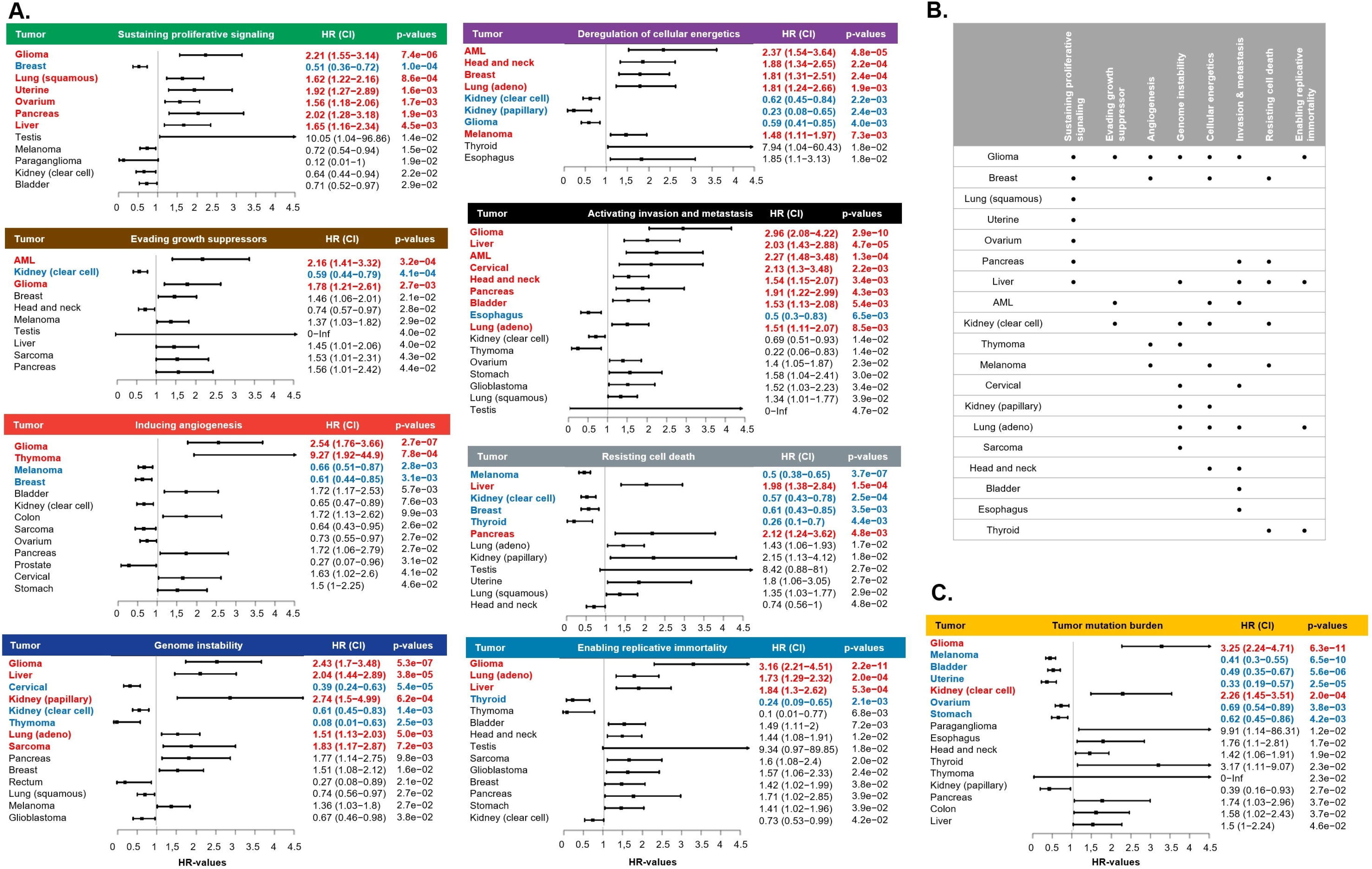
Effect of hallmark signatures **(A)** and tumor mutation burden **(C)** on patient survival. Summary of the significant prognostic hallmark signatures in different types of tumors **(B)**.

Of the eight hallmark feature signatures, seven showed a significant association with OS in low grade glioma. On the other hand, lung squamous carcinoma, uterine, ovarian, sarcoma, bladder and esophageal cancer contained only one significant hallmark signature **(Figure 3B)**.

Tumor mutation burden was also determined, and it showed a significant association with OS in glioma (HR=3.25, *P*=6.3E-11), melanoma (HR=0.41, *P*=6.5E-10), bladder cancer (HR=0.49, *P*=5.6E-06), uterine cancer (HR=0.33, *P*=2.5E-05), ovarian cancer (HR=0.69, *P*=3.8E-03), stomach cancer (HR=0.62, *P*=4.2E-03) and kidney renal clear cell carcinoma (HR=2.26, *P*=2.0E-04) **(Figure 3C)**.

In multivariate analysis of OS, including the expression signature of hallmark features, sex, race, tumor stage, tumor grade and age, most of the signatures retained their significance **(Table 2)**.

**Table 2.** Multivariate Cox regression analysis of hallmark gene signatures after including sex, race, stage, grade and age. Significant P (P<0.05) and HR values in univariate and both uni- and multivariate survival analyses are green and red, respectively. HR values with asterisk (*) shows that there are not any events in one of the groups in the survival analysis. *

|  | Sustaining proliferative signaling |  | Resisting cell death |  | Inducing angiogenesis |  | Genome instability |  | Evading growth suppressors |  | Enabling replicative immortality |  | Deregulation of cellular energetics |  | Activation invasion and metastasis |  |
| --- | --- | --- | --- | --- | --- | --- | --- | --- | --- | --- | --- | --- | --- | --- | --- | --- |
| Tumor types | P | HR | P | HR | P | HR | P | HR | P | HR | P | HR | P | HR | P | HR |
| Bladder | 9.90E-09 | 0.78 | 1.45E-08 | 0.8 | 8.23E-09 | 1.48 | 1.92E-08 | 0.86 | 1.95E-08 | 0.86 | 5.56E-09 | 1.4 | 1.61E-08 | 1.17 | 6.76E-09 | 1.37 |
| Breast | 1.05E-16 | 0.64 | 8.41E-17 | 0.69 | 3.23E-16 | 0.73 | 1.67E-16 | 1.57 | 1.59E-16 | 1.42 | 7.19E-18 | 1.88 | 1.93E-17 | 1.59 | 4.02E-16 | 1.34 |
| Cervical | n.s. | 0.82 | n.s. | 1.08 | 4.85E-02 | 1.73 | 7.82E-05 | 0.32 | n.s. | 1.25 | n.s. | 1.3 | n.s. | 0.81 | 1.14E-02 | 2.19 |
| Colon | 1.45E-05 | 1.02 | 1.93E-06 | 0.55 | 1.31E-05 | 1.2 | 1.36E-05 | 0.97 | 1.29E-06 | 0.51 | 5.66E-06 | 1.57 | 1.40E-05 | 0.97 | 1.44E-05 | 1.01 |
| Esophagus | 1.94E-02 | 0.84 | 1.73E-02 | 0.72 | 1.72E-02 | 0.77 | 1.77E-02 | 1.21 | 2.01E-02 | 0.93 | 9.40E-03 | 2.16 | 2.60E-04 | 3.68 | 1.80E-02 | 0.77 |
| Glioblastoma | 1.38E-03 | 1.62 | 1.91E-03 | 1.53 | 7.66E-03 | 1.22 | 1.09E-03 | 0.64 | 2.44E-03 | 0.68 | 1.78E-03 | 1.51 | 8.59E-03 | 1.18 | 7.36E-03 | 1.26 |
| Head and neck | 3.24E-05 | 0.81 | 5.94E-05 | 0.87 | 1.74E-05 | 1.34 | 2.89E-05 | 1.28 | 4.71E-05 | 0.85 | 4.79E-05 | 1.17 | 1.72E-06 | 1.83 | 6.61E-06 | 1.49 |
| Kidney (clear cell) | 1.60E-24 | 0.85 | 1.77E-25 | 0.69 | 8.43E-25 | 0.86 | 1.08E-25 | 0.69 | 3.02E-25 | 0.73 | 1.25E-24 | 0.86 | 6.68E-26 | 0.67 | 6.87E-25 | 0.78 |
| Kidney (papillary) | 4.69E-10 | 2.8 | 6.04E-10 | 2.76 | 5.53E-09 | 0.54 | 3.38E-09 | 2.04 | 3.08E-09 | 2.64 | 1.84E-09 | 2.29 | 5.41E-12 | 0.06 | 7.56E-09 | 1.49 |
| AML | 8.22E-07 | 0.62 | 2.75E-06 | 0.76 | 4.57E-06 | 1.15 | 3.29E-06 | 1.28 | 1.44E-07 | 1.78 | 1.67E-06 | 1.41 | 6.19E-10 | 2.69 | 4.58E-08 | 1.98 |
| Glioma | 5.29E-21 | 1.82 | 7.40E-19 | 0.91 | 5.72E-22 | 2.12 | 1.26E-20 | 1.7 | 2.49E-19 | 1.35 | 9.92E-24 | 2.28 | 9.58E-22 | 0.5 | 2.48E-24 | 2.67 |
| Liver | 1.09E-05 | 1.57 | 2.40E-06 | 1.86 | 3.66E-05 | 0.7 | 1.01E-06 | 1.94 | 3.02E-05 | 1.37 | 2.89E-06 | 1.72 | 8.93E-05 | 1.09 | 1.12E-06 | 1.86 |
| Lung (adeno) | 8.35E-08 | 1.36 | 1.35E-07 | 1.26 | 1.73E-07 | 0.84 | 1.22E-08 | 1.53 | 1.29E-07 | 1.31 | 4.11E-09 | 1.65 | 6.27E-08 | 1.53 | 5.86E-08 | 1.43 |
| Lung (squamous) | 8.48E-07 | 1.99 | 9.11E-05 | 1.45 | 3.73E-04 | 1.34 | 1.54E-04 | 0.71 | 2.09E-04 | 0.71 | 1.09E-03 | 1.1 | 7.24E-04 | 0.83 | 2.79E-04 | 1.34 |
| Ovary | 1.68E-04 | 1.53 | 4.45E-03 | 0.87 | 1.05E-03 | 0.75 | 1.88E-03 | 0.77 | 5.94E-03 | 1.08 | 3.14E-03 | 0.83 | 4.26E-03 | 0.85 | 1.14E-03 | 1.36 |
| Pancreas | 7.58E-03 | 2.03 | 3.70E-02 | 1.82 | n.s. | 1.51 | 4.84E-02 | 1.52 | n.s. | 1.37 | n.s. | 1.42 | n.s. | 1.32 | 1.53E-02 | 1.81 |
| Paraganglioma | 6.27E-02 | 0.12 | n.s. | 3.61 | n.s. | 0.25 | n.s. | 4.57 | n.s. | 2.73 | n.s. | 1.69 | n.s. | * | n.s. | 0.48 |
| Prostate | n.s. | * | n.s. | inf | 9.98E-02 | * | n.s. | inf | n.s. | * | n.s. | * | n.s. | inf | n.s. | * |
| Rectum | 1.77E-02 | 2.8 | 1.36E-02 | 0.49 | 8.56E-03 | 0.44 | 2.90E-02 | 0.6 | 2.24E-02 | 0.64 | 3.54E-02 | 1.02 | 3.28E-02 | 1.39 | 3.53E-02 | 1.23 |
| Sarcoma | 2.83E-02 | 1.51 | n.s. | 0.73 | 2.47E-03 | 0.53 | 2.73E-03 | 2.01 | 2.40E-02 | 1.49 | 2.56E-02 | 1.47 | n.s. | 1.18 | n.s. | 0.71 |
| Melanoma | 4.35E-10 | 0.67 | 4.29E-13 | 0.5 | 1.12E-10 | 0.61 | 8.21E-11 | 1.63 | 9.88E-09 | 1.1 | 2.58E-09 | 0.75 | 1.63E-10 | 1.6 | 9.99E-09 | 0.93 |
| Stomach | 2.15E-03 | 1.14 | 2.20E-03 | 1.19 | 1.42E-03 | 1.35 | 1.28E-03 | 0.75 | 3.74E-04 | 0.64 | 1.67E-03 | 1.21 | 2.50E-03 | 0.92 | 1.00E-03 | 1.48 |
| Testis | 5.88E-03 | * | 5.72E-03 | * | 3.58E-03 | * | 2.96E-03 | >100 | 4.93E-03 | * | 5.81E-03 | * | 5.87E-03 | >100 | 4.56E-03 | * |
| Thyroid | 1.73E-10 | 0.4 | 6.54E-11 | 0.34 | 1.52E-11 | 3.38 | 2.36E-10 | 0.77 | 6.82E-11 | 2.02 | 6.40E-13 | 0.35 | 1.31E-11 | 6.24 | 2.29E-10 | 0.59 |
| Thymoma | n.s. | 0.43 | n.s. | 2.35 | 1.24E-02 | 7.68 | 1.65E-02 | 0.08 | n.s. | 0.25 | 8.35E-03 | 0.04 | 4.97E-02 | 4.11 | 2.83E-02 | 0.2 |
| Uterine | 2.07E-07 | 1.56 | 9.32E-07 | 1.54 | 1.34E-06 | 0.85 | 1.58E-06 | 1.21 | 7.64E-07 | 1.43 | 1.01E-06 | 1.32 | 1.89E-06 | 1.02 | 1.62E-06 | 0.82 |

### Genes with the greatest prognostic power in multiple tumor types

In at least ten tumor types, there were 39 genes whose expression was associated with OS **(Figure 4A)**. We pinpointed the genes with the highest prognostic power in each cancer hallmark feature: BRCA1 associated with genome instability in low grade glioma (HR=4.26, *P*<1E-16), CDK1 linked to cell death resistance in kidney papillary carcinoma (HR=5.67, *P*=2.1E-10), the E2F1 tumor suppressor in cervical cancer (HR=0.38, *P*=2.4E-05), EREG enabling replicative immortality in cervical cancer (HR=3.23, *P*=2.1E-07), FBP1 participating in the deregulation of cellular energetics in kidney renal clear cell carcinoma (HR=0.45, *P*=2.8E-07), MYC activating invasion and metastasis in bladder cancer (HR=1.81, *P*=5.8E-05), RUNX1 sustaining proliferative signaling in glioma (HR=2.96, *P*=3.1E-10) and SERPINE1 playing a role in inducing angiogenesis in glioma (HR=3.36, *P*=1.5E-12) **(Figure 4B-I)**.

**Figure 4.**
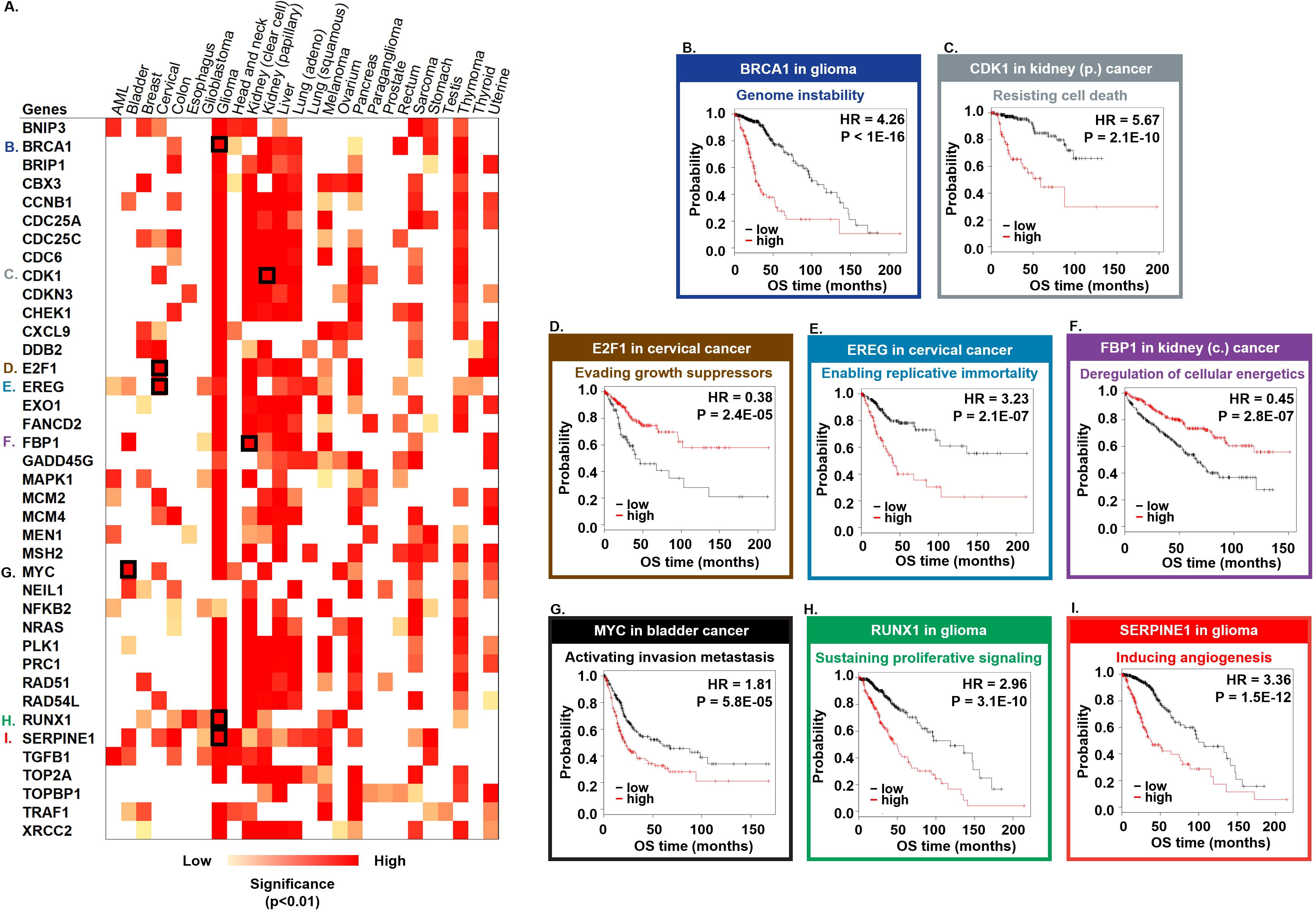
Best performing genes in at least 10 distinct tumor types.

In addition, multivariate Cox regression analysis was also performed using the expression of the 39 most significant genes and the available clinical variables, including race, sex, age, tumor stage and tumor grade. Of the clinical parameters, age and tumor stage were the variables that reached significance in the Cox model in most tumors ( for detailed results, see **Supplemental Table 2)**.

## DISCUSSION

In this study, we examined the prognostic significance of previously established cancer hallmark genes ^5^. For the survival analysis, we utilized an RNA-seq database from the TCGA that contains 9,720 patients of 26 tumor types with clinical annotations. Kidney renal clear cell carcinoma, low grade glioma and melanoma had the highest proportion of cancer hallmark genes that correlated with survival. Hierarchical clustering analysis showed that some cancer hallmark genes clustered together, such as those involved with invasion and metastasis activation, genome instability, sustained proliferative signaling and cellular energetics deregulation (distance was based on the percentage of significant genes per hallmark in each tumor type).

A transcriptomic surrogate signature for each hallmark was also determined; this is based on the means of the average expression of the cancer genes associated with the given hallmark. The prognostic significance of these factors was examined in different types of cancers. Among the eight main hallmark signatures, those associated with oncogene activation, genome instability, cellular energetics, invasion and metastasis and cell death resistance were significant in at least five tumor types.

Oncogenes have a major role in the control of cell proliferation, differentiation and survival during tumorigenesis. c-MYC was the first characterized oncogene that is activated by chromosome translocation in human Burkitt’s lymphomas ^15^. Expression of the altered c-MYC gene is increased in tumor cells and is associated with extensive cell proliferation and contributes to tumor development. The association between c-MYC expression and patient survival remains controversial ^15^, and we observed a worse prognosis in patients with higher expression of c-MYC. Similar results were present in the case of the ERBB2 gene, which encodes a cell surface protein-tyrosine kinase receptor that is associated with the progression of breast cancer ^16^ and higher expression of genes in the Wnt-β-catenin pathway. This pathway is mutated in more than 85% of colorectal cancers ^17^. β-catenin (CTNNB1) is the most frequently mutated gene, and it can be detected in more than 80% of colorectal tumors. In addition, high expression of CTNNB1 is associated with shorter survival in colorectal cancer ^17^. Finally, overexpression of cyclin D1 (CCND1), a member of the cyclin family, also correlated with poor survival in esophageal squamous cell carcinoma ^18^.

Chromosomal instability (CIN) and microsatellite instability (MSI) are the two main types of genomic instability in human cancers ^4^. The expression of genomic instability-related genes is higher in metastatic samples than in primary tumors ^19^. In breast cancer, Habermann *et al* performed gene expression profiling in which they examined the correlation between gene expression, genome instability and clinical outcomes ^20^ and identified a 12‐gene aneuploidy‐specific signature that is an independent predictor of clinical outcome. In our analysis, the transcriptomic signature consisting of 150 genes contributing to genome instability ^5^ was prognostic in eight tumors. Among these, high signature expression was associated with poor survival in low grade glioma, liver cancer, kidney papillary cancer, lung adenocarcinoma and sarcoma. In cervical cancer, renal clear cell carcinoma and thymoma, the high expression of the hallmark signature was correlated with a favorable outcome.

Altered energy metabolism involves an increased rate of glycolysis and limited oxidative phosphorylation. These features of proliferating cancer cells enable the retention of macromolecules, which help to drive constitutive cell growth and proliferation ^4^. Among the numerous metabolic pathway-associated genes, the high expression of GLUT1, G6PD, TKTL1 and PGI/AMF are significantly correlated with decreased survival in breast cancer ^21^. The FAS gene is upregulated at an early stage in multiple cancers, including breast ^22^, stomach ^23^ and prostate cancers ^24^; its expression is positively correlated with poor survival. Our results show that the high expression of the transcriptomic signature of cancer metabolism-associated genes is linked to decreased survival in acute myeloid leukemia, head and neck cancers, breast cancer, lung adenocarcinoma and melanoma. However, in kidney renal clear cell carcinoma, kidney papillary cancer and low grade glioma, the high expression of the signature was associated with a better outcome.

Epithelial-mesenchymal transition (EMT) is a multistep process that contributes to the migratory and invasive capacity of cells, which are essential for the development and metastasis of cancer ^4^. In many types of cancer, including breast and head and neck cancers, developmental EMT pathways such as Notch have been reported to be dysregulated, and activation of these pathways often correlates with poor survival ^25^. The suppression of EMT results in the increase of cell proliferation with increased expression of nucleoside transporters in pancreatic tumors. These changes lead to enhanced sensitivity to gemcitabine treatment and increased overall survival in mice ^26^. The importance of EMT is supported by our observation that the transcriptomic signature of the tumor invasion and metastasis activation-associated genes ^5^ had prognostic significance in the highest number of tumors. Among the tumors, the high expression of the signature was linked to poor survival outcome in low grade glioma, liver cancer, acute myeloid leukemia, cervical cancer, head and neck cancers, pancreas cancer, bladder cancer and lung adenocarcinoma.

The resistance of cancer cells to apoptosis is a fundamental aspect of cancer development, which includes the upregulation of antiapoptotic proteins and the downregulation of proapoptotic proteins ^27^. The number of gene expression signature studies of apoptotic genes is limited, and studies more commonly reflect on single apoptotic genes. Holleman *et al* performed a microarray gene expression study in which they examined the expression pattern of 70 key apoptotic genes in acute lymphoblastic leukemia (ALL) and concluded that leukemia subtypes have a unique expression pattern of apoptosis genes and that select genes are linked to cellular drug resistance and prognosis in childhood B-lineage ALL ^28^. Another study investigated 40 genes involved in the extrinsic and intrinsic pathways in myeloma cells, and these genes were linked to poor prognosis and were overexpressed in normal plasmablastic cells ^29^. In our study, the cell death resistance signature based on a set of 119 genes ^30 31^ was linked to poor survival in liver and pancreatic cancers and good survival in melanoma, kidney renal clear cell carcinoma, breast cancer and thyroid cancer.

In brief, RNA-seq-based transcriptomic data were utilized to perform survival analysis across 26 different types of cancer. Strikingly, the signatures constructed from the cancer hallmark genes showed tumor type-specific correlations with survival. Individual cancer hallmark genes showing prognostic significance in more than 10 cancer types were also uncovered. These results help to prioritize targeting the most relevant hallmark for drug development in each tumor type.

## ACKNOWLEDGEMENTS

The research was financed by the 2018-2.1.17-TET-KR-00001 and KH-129581 grants and by the Higher Education Institutional Excellence Programme of the Ministry for Innovation and Technology in Hungary, within the framework of the Bionic thematic programme of the Semmelweis University. This study was also supported by the ÚNKP-19-3-IV-SE-5 New National Excellence Program of the Ministry for Innovation and Technology. The authors acknowledge the support of ELIXIR Hungary (www.elixir-hungary.org).

## AUTHORS’ CONTRIBUTIONS

B.G. contributed to the conception, design and writing of the manuscript. G.M. contributed to the data interpretation and drafting the manuscript. Á.N. contributed to the data analysis, data interpretation and drafting the manuscript. All of the authors read and approved the final manuscript.

## DATA ACCESSIBILITY

TCGA (The Cancer Genome Atlas) dataset is available using the following link: https://portal.gdc.cancer.gov/

## SUPPLEMENTARY MATERIALS

**Supplemental Table 1.** Univariate analysis of cancer hallmark genes across 26 types of cancer.

**Supplemental Table 2.** Multivariate analysis of genes that were significant in more than 10 tumors.

