## Supplemental Tables for "Pancancer survival analysis of cancer hallmark genes": Supplemental_Table_2.docx

**Supplemental Table 2.** Multivariate analysis of genes that were significant in more than 10 tumor types.

|  | **Bladder** | | | | | | | | | | | | | |
| --- | --- | --- | --- | --- | --- | --- | --- | --- | --- | --- | --- | --- | --- | --- |
|  | **Univariate** | | **Multivariate** | | | | | | | | | | | |
|  | **n=405** | | **n=405** | | **n=405** | | **n=388** | | **n=405** | | **n=403** | | **n=402** | |
| **Gene** | **Expression P-value** | **Expression HR** | **Expression P-value** | **Expression HR** | **Gender P-value** | **Gender HR** | **Race P-value** | **Race HR** | **Age P-value** | **Age HR** | **Stage P-value** | **Stage HR** | **Grade P-value** | **Grade HR** |
| **MYC** | 1.65E-04 | 1.75 | 4.00E-04 | 1.75 | 7.06E-01 | 0.94 | 2.76E-01 | 1.16 | 2.46E-04 | 1.03 | 3.30E-07 | 1.68 | 8.68E-01 | 1.13 |
| **FBP1** | 1.61E-05 | 0.52 | 9.00E-04 | 0.59 | 5.90E-01 | 0.91 | 2.52E-01 | 1.17 | 2.49E-04 | 1.03 | 8.76E-07 | 1.65 | 6.98E-01 | 1.33 |
| **TGFB1** | 3.42E-03 | 1.61 | 1.00E-03 | 1.74 | 4.72E-01 | 0.88 | 1.71E-01 | 1.21 | 2.24E-04 | 1.03 | 8.45E-08 | 1.74 | 5.84E-01 | 1.49 |
| **EREG** | 6.15E-03 | 1.61 | 6.40E-03 | 1.64 | 4.31E-01 | 0.87 | 1.85E-01 | 1.2 | 1.75E-04 | 1.03 | 2.94E-07 | 1.68 | 6.58E-01 | 1.38 |
| **SERPINE1** | 1.18E-03 | 1.67 | 9.30E-03 | 1.54 | 5.29E-01 | 0.9 | 2.24E-01 | 1.19 | 2.21E-04 | 1.03 | 1.27E-06 | 1.64 | 7.35E-01 | 1.28 |
| **CCNB1** | 4.32E-03 | 1.54 | 1.08E-02 | 1.5 | 5.21E-01 | 0.89 | 3.39E-01 | 1.14 | 2.19E-04 | 1.03 | 2.02E-07 | 1.69 | 8.44E-01 | 1.16 |
| **TRAF1** | 7.98E-03 | 0.61 | 2.30E-02 | 0.65 | 3.25E-01 | 0.84 | 2.71E-01 | 1.17 | 7.88E-04 | 1.03 | 4.94E-07 | 1.66 | 6.27E-01 | 1.43 |
| **PLK1** | 8.68E-03 | 1.49 | 2.74E-02 | 1.42 | 4.60E-01 | 0.88 | 3.44E-01 | 1.14 | 1.85E-04 | 1.03 | 3.77E-07 | 1.67 | 7.96E-01 | 1.21 |
| **NEIL1** | 2.02E-03 | 0.59 | 2.85E-02 | 0.67 | 6.49E-01 | 0.92 | 2.45E-01 | 1.18 | 3.74E-04 | 1.03 | 7.90E-07 | 1.65 | 7.82E-01 | 1.22 |
| **CDKN3** | 7.06E-02 | 1.32 | 2.89E-02 | 1.41 | 4.65E-01 | 0.88 | 3.32E-01 | 1.14 | 1.95E-04 | 1.03 | 1.74E-07 | 1.7 | 7.71E-01 | 1.24 |
|  | **Breast** | | | | | | | | | | | | | |
|  | **Univariate** | | **Multivariate** | | | | | | | | | | | |
|  | **n=1090** | | **n=1090** | | **n=1090** | | **n=995** | | **n=1090** | | **n=1067** | | **n=0** | |
| **Gene** | **Expression P-value** | **Expression HR** | **Expression P-value** | **Expression HR** | **Gender P-value** | **Gender HR** | **Race P-value** | **Race HR** | **Age P-value** | **Age HR** | **Stage P-value** | **Stage HR** | **Grade P-value** | **Grade HR** |
| **DDB2** | 8.17E-04 | 0.57 | 1.00E-04 | 0.49 | 5.02E-01 | 0.51 | 2.38E-01 | 1.14 | 6.64E-08 | 1.04 | 2.01E-14 | 2.51 | No data | No data |
| **FBP1** | 4.18E-02 | 0.72 | 1.00E-04 | 0.49 | 5.15E-01 | 0.52 | 4.38E-01 | 1.09 | 1.34E-09 | 1.04 | 2.07E-13 | 2.38 | No data | No data |
| **XRCC2** | 9.97E-03 | 1.54 | 1.00E-04 | 2.11 | 3.57E-01 | 0.39 | 4.73E-01 | 1.08 | 1.52E-08 | 1.04 | 6.73E-13 | 2.39 | No data | No data |
| **CBX3** | 4.02E-04 | 1.75 | 3.00E-04 | 1.89 | 3.70E-01 | 0.4 | 3.53E-01 | 1.11 | 2.16E-07 | 1.04 | 2.01E-12 | 2.28 | No data | No data |
| **RAD51** | 1.86E-03 | 1.67 | 3.00E-04 | 1.94 | 3.42E-01 | 0.38 | 6.13E-01 | 1.06 | 1.21E-08 | 1.04 | 1.24E-12 | 2.34 | No data | No data |
| **MCM4** | 5.30E-02 | 1.41 | 7.00E-04 | 1.94 | 3.77E-01 | 0.41 | 3.15E-01 | 1.12 | 3.80E-09 | 1.04 | 8.43E-14 | 2.45 | No data | No data |
| **CDC25C** | 2.82E-03 | 1.64 | 1.00E-03 | 1.81 | 3.38E-01 | 0.38 | 2.80E-01 | 1.12 | 2.39E-08 | 1.04 | 2.85E-12 | 2.28 | No data | No data |
| **RAD54L** | 4.18E-02 | 1.39 | 1.40E-03 | 1.78 | 3.81E-01 | 0.41 | 5.30E-01 | 1.07 | 3.72E-09 | 1.04 | 9.89E-13 | 2.31 | No data | No data |
| **CHEK1** | 7.38E-02 | 1.33 | 3.00E-03 | 1.69 | 3.74E-01 | 0.41 | 3.24E-01 | 1.11 | 1.64E-08 | 1.04 | 5.34E-13 | 2.33 | No data | No data |
| **TGFB1** | 1.54E-02 | 0.57 | 3.00E-03 | 0.48 | 4.53E-01 | 0.47 | 1.54E-01 | 1.17 | 5.20E-08 | 1.04 | 1.94E-13 | 2.38 | No data | No data |
|  | **Cervical** | | | | | | | | | | | | | |
|  | **Univariate** | | **Multivariate** | | | | | | | | | | | |
|  | **n=304** | | **n=304** | | **n=304** | | **n=259** | | **n=304** | | **n=0** | | **n=272** | |
| **Gene** | **Expression P-value** | **Expression HR** | **Expression P-value** | **Expression HR** | **Gender P-value** | **Gender HR** | **Race P-value** | **Race HR** | **Age P-value** | **Age HR** | **Stage P-value** | **Stage HR** | **Grade P-value** | **Grade HR** |
| **CDK1** | 1.62E-03 | 0.46 | 1.00E-04 | 0.34 | - | - | 9.01E-01 | 0.98 | 8.89E-03 | 1.03 | No data | No data | 8.31E-01 | 1.06 |
| **EREG** | 3.25E-07 | 3.23 | 1.00E-04 | 2.86 | - | - | 7.75E-01 | 0.95 | 2.43E-02 | 1.02 | No data | No data | 5.01E-01 | 1.21 |
| **E2F1** | 7.28E-05 | 0.40 | 1.40E-03 | 0.42 | - | - | 9.72E-01 | 1.01 | 2.04E-02 | 1.02 | No data | No data | 5.27E-01 | 1.19 |
| **CDC25C** | 5.99E-03 | 0.52 | 2.00E-03 | 0.44 | - | - | 7.38E-01 | 0.94 | 1.75E-02 | 1.02 | No data | No data | 8.35E-01 | 1.06 |
| **RAD54L** | 2.14E-03 | 0.48 | 2.00E-03 | 0.43 | - | - | 7.13E-01 | 0.93 | 2.41E-02 | 1.02 | No data | No data | 7.56E-01 | 1.09 |
| **MYC** | 3.12E-03 | 2.04 | 5.30E-03 | 2.22 | - | - | 9.14E-01 | 0.98 | 6.85E-03 | 1.03 | No data | No data | 9.42E-01 | 0.98 |
| **CDC25A** | 1.12E-01 | 0.68 | 5.40E-03 | 0.46 | - | - | 6.54E-01 | 0.92 | 7.59E-03 | 1.03 | No data | No data | 7.69E-01 | 1.09 |
| **DDB2** | 2.11E-04 | 0.42 | 6.30E-03 | 0.48 | - | - | 8.05E-01 | 0.95 | 3.13E-02 | 1.02 | No data | No data | 8.42E-01 | 1.06 |
| **MCM4** | 3.92E-03 | 0.46 | 6.80E-03 | 0.42 | - | - | 8.27E-01 | 0.96 | 2.98E-02 | 1.02 | No data | No data | 5.66E-01 | 1.17 |
| **RUNX1** | 1.93E-02 | 1.79 | 8.30E-03 | 2.08 | - | - | 9.98E-01 | 1 | 2.17E-02 | 1.02 | No data | No data | 9.91E-01 | 1 |
|  | **Colon** | | | | | | | | | | | | | |
|  | **Univariate** | | **Multivariate** | | | | | | | | | | | |
|  | **n=454** | | **n=454** | | **n=454** | | **n=282** | | **n=454** | | **n=443** | | **n=0** | |
| **Gene** | **Expression P-value** | **Expression HR** | **Expression P-value** | **Expression HR** | **Gender P-value** | **Gender HR** | **Race P-value** | **Race HR** | **Age P-value** | **Age HR** | **Stage P-value** | **Stage HR** | **Grade P-value** | **Grade HR** |
| **EREG** | 1.07E-02 | 0.58 | 2.40E-03 | 0.43 | 6.38E-01 | 1.13 | 5.40E-01 | 1.1 | 2.32E-02 | 1.02 | 7.29E-08 | 2.37 | No data | No data |
| **CCNB1** | 3.26E-03 | 0.47 | 2.09E-02 | 0.41 | 4.80E-01 | 1.2 | 7.06E-01 | 1.06 | 1.44E-02 | 1.03 | 5.25E-06 | 2.05 | No data | No data |
| **CBX3** | 6.05E-02 | 1.47 | 2.76E-02 | 1.78 | 9.28E-01 | 1.02 | 4.05E-01 | 1.14 | 1.22E-02 | 1.03 | 2.20E-06 | 2.11 | No data | No data |
| **MEN1** | 4.25E-02 | 1.54 | 3.05E-02 | 1.8 | 5.18E-01 | 1.18 | 4.79E-01 | 1.12 | 3.23E-03 | 1.03 | 9.49E-07 | 2.16 | No data | No data |
| **NEIL1** | 3.73E-03 | 1.89 | 3.91E-02 | 1.74 | 6.11E-01 | 1.14 | 7.36E-01 | 1.05 | 5.89E-03 | 1.03 | 6.02E-07 | 2.2 | No data | No data |
| **XRCC2** | 2.34E-01 | 1.28 | 4.20E-02 | 1.69 | 6.85E-01 | 1.11 | 5.46E-01 | 1.1 | 7.52E-03 | 1.03 | 8.84E-07 | 2.17 | No data | No data |
|  | **Esophagus** | | | | | | | | | | | | | |
|  | **Univariate** | | **Multivariate** | | | | | | | | | | | |
|  | **n=161** | | **n=161** | | **n=161** | | **n=143** | | **n=161** | | **n=142** | | **n=126** | |
| **Gene** | **Expression P-value** | **Expression HR** | **Expression P-value** | **Expression HR** | **Gender P-value** | **Gender HR** | **Race P-value** | **Race HR** | **Age P-value** | **Age HR** | **Stage P-value** | **Stage HR** | **Grade P-value** | **Grade HR** |
| **RUNX1** | 1.03E-03 | 0.44 | 1.17E-02 | 0.37 | 6.08E-02 | 7.37 | 5.48E-01 | 1.26 | 3.66E-01 | 1.02 | 1.43E-02 | 2.27 | 4.20E-01 | 0.72 |
| **BRIP1** | 2.37E-01 | 0.72 | 4.17E-02 | 0.34 | 1.45E-01 | 4.57 | 4.09E-01 | 1.38 | 9.20E-01 | 1 | 1.70E-02 | 2.26 | 3.19E-01 | 0.66 |
|  | **Glioblastoma** | | | | | | | | | | | | | |
|  | **Univariate** | | **Multivariate** | | | | | | | | | | | |
|  | **n=153** | | **n=153** | | **n=153** | | **n=152** | | **n=153** | | **n=0** | | **n=0** | |
| **Gene** | **Expression P-value** | **Expression HR** | **Expression P-value** | **Expression HR** | **Gender P-value** | **Gender HR** | **Race P-value** | **Race HR** | **Age P-value** | **Age HR** | **Stage P-value** | **Stage HR** | **Grade P-value** | **Grade HR** |
| **FBP1** | 9.21E-03 | 1.64 | 4.70E-03 | 1.71 | 6.73E-01 | 0.92 | 1.18E-01 | 1.38 | 5.62E-04 | 1.03 | No data | No data | No data | No data |
| **MAPK1** | 7.24E-03 | 0.60 | 7.50E-03 | 0.59 | 7.07E-01 | 0.93 | 1.15E-01 | 1.39 | 8.72E-04 | 1.03 | No data | No data | No data | No data |
| **RUNX1** | 5.36E-03 | 1.75 | 8.60E-03 | 1.74 | 9.84E-01 | 1 | 2.28E-01 | 1.29 | 7.35E-04 | 1.03 | No data | No data | No data | No data |
| **EREG** | 4.79E-03 | 1.79 | 1.34E-02 | 1.69 | 9.03E-01 | 0.98 | 1.90E-01 | 1.31 | 1.12E-03 | 1.02 | No data | No data | No data | No data |
| **SERPINE1** | 8.04E-03 | 1.75 | 1.51E-02 | 1.72 | 4.24E-01 | 0.85 | 1.62E-01 | 1.33 | 1.39E-03 | 1.02 | No data | No data | No data | No data |
| **TGFB1** | 4.34E-03 | 1.72 | 2.13E-02 | 1.63 | 6.04E-01 | 0.9 | 1.76E-01 | 1.32 | 2.30E-03 | 1.02 | No data | No data | No data | No data |
| **NFKB2** | 6.41E-03 | 1.72 | 2.28E-02 | 1.6 | 9.35E-01 | 1.02 | 1.50E-01 | 1.34 | 1.65E-03 | 1.02 | No data | No data | No data | No data |
| **CDK1** | 5.51E-02 | 0.70 | 2.59E-02 | 0.66 | 5.55E-01 | 0.89 | 1.13E-01 | 1.39 | 4.86E-04 | 1.03 | No data | No data | No data | No data |
| **NEIL1** | 2.52E-02 | 1.52 | 3.57E-02 | 1.48 | 9.62E-01 | 1.01 | 1.50E-01 | 1.35 | 5.67E-04 | 1.03 | No data | No data | No data | No data |
| **NRAS** | 1.49E-02 | 0.63 | 4.73E-02 | 0.67 | 5.28E-01 | 0.88 | 1.93E-01 | 1.31 | 2.29E-03 | 1.02 | No data | No data | No data | No data |
|  | **Head and neck** | | | | | | | | | | | | | |
|  | **Univariate** | | **Multivariate** | | | | | | | | | | | |
|  | **n=500** | | **n=500** | | **n=500** | | **n=483** | | **n=500** | | **n=432** | | **n=481** | |
| **Gene** | **Expression P-value** | **Expression HR** | **Expression P-value** | **Expression HR** | **Gender P-value** | **Gender HR** | **Race P-value** | **Race HR** | **Age P-value** | **Age HR** | **Stage P-value** | **Stage HR** | **Grade P-value** | **Grade HR** |
| **BNIP3** | 1.73E-03 | 1.54 | 2.10E-03 | 1.62 | 2.16E-01 | 0.81 | 1.64E-01 | 1.19 | 7.61E-04 | 1.03 | 6.73E-05 | 1.48 | 5.07E-01 | 1.13 |
| **SERPINE1** | 1.10E-04 | 1.85 | 2.90E-03 | 1.75 | 3.12E-01 | 0.84 | 1.08E-01 | 1.22 | 8.54E-04 | 1.03 | 1.77E-04 | 1.44 | 7.56E-01 | 1.06 |
| **CDC6** | 3.04E-02 | 1.37 | 7.90E-03 | 1.57 | 9.64E-02 | 0.75 | 2.64E-01 | 1.15 | 1.92E-03 | 1.02 | 1.29E-04 | 1.45 | 9.30E-01 | 0.98 |
| **PLK1** | 4.65E-02 | 1.32 | 1.23E-02 | 1.49 | 1.51E-01 | 0.78 | 9.36E-02 | 1.23 | 1.57E-03 | 1.02 | 1.39E-04 | 1.45 | 8.66E-01 | 0.97 |
| **TRAF1** | 3.16E-03 | 0.67 | 1.28E-02 | 0.68 | 1.71E-01 | 0.79 | 1.41E-01 | 1.2 | 1.66E-03 | 1.03 | 8.08E-05 | 1.48 | 6.82E-01 | 1.08 |
| **TGFB1** | 4.97E-04 | 1.82 | 1.39E-02 | 1.63 | 2.49E-01 | 0.82 | 1.72E-01 | 1.18 | 1.38E-03 | 1.03 | 9.27E-05 | 1.46 | 6.23E-01 | 1.09 |
| **CBX3** | 9.57E-03 | 1.47 | 1.90E-02 | 1.49 | 1.70E-01 | 0.79 | 1.27E-01 | 1.21 | 1.08E-03 | 1.03 | 9.76E-05 | 1.47 | 9.29E-01 | 0.98 |
| **MYC** | 4.27E-03 | 1.56 | 2.50E-02 | 1.5 | 2.53E-01 | 0.82 | 2.12E-01 | 1.17 | 2.19E-03 | 1.02 | 7.61E-05 | 1.47 | 6.52E-01 | 1.08 |
| **CHEK1** | 1.34E-01 | 1.25 | 2.68E-02 | 1.45 | 1.63E-01 | 0.79 | 2.25E-01 | 1.16 | 1.55E-03 | 1.03 | 7.92E-05 | 1.47 | 9.82E-01 | 1 |
| **GADD45G** | 1.72E-01 | 0.82 | 3.22E-02 | 0.7 | 3.10E-01 | 0.84 | 1.23E-01 | 1.21 | 9.10E-04 | 1.03 | 6.59E-05 | 1.48 | 6.92E-01 | 1.07 |
|  | **Kidney (clear cell)** | | | | | | | | | | | | | |
|  | **Univariate** | | **Multivariate** | | | | | | | | | | | |
|  | **n=530** | | **n=530** | | **n=530** | | **n=523** | | **n=530** | | **n=527** | | **n=522** | |
| **Gene** | **Expression P-value** | **Expression HR** | **Expression P-value** | **Expression HR** | **Gender P-value** | **Gender HR** | **Race P-value** | **Race HR** | **Age P-value** | **Age HR** | **Stage P-value** | **Stage HR** | **Grade P-value** | **Grade HR** |
| **MAPK1** | 2.22E-07 | 0.43 | 0.00E+00 | 0.46 | 3.82E-01 | 0.87 | 7.31E-01 | 0.95 | 2.60E-05 | 1.03 | 8.59E-14 | 1.73 | 2.17E-02 | 1.55 |
| **RUNX1** | 1.04E-09 | 2.50 | 0.00E+00 | 2.15 | 3.00E-01 | 0.84 | 9.39E-01 | 0.99 | 1.95E-05 | 1.03 | 1.39E-14 | 1.76 | 4.21E-02 | 1.48 |
| **FBP1** | 4.87E-07 | 0.46 | 1.00E-04 | 0.51 | 2.47E-01 | 0.82 | 8.27E-01 | 1.04 | 4.19E-05 | 1.03 | 4.07E-14 | 1.74 | 3.30E-02 | 1.5 |
| **NEIL1** | 2.41E-03 | 1.61 | 2.00E-04 | 1.82 | 8.95E-01 | 1.02 | 7.70E-01 | 0.96 | 3.89E-05 | 1.03 | 8.66E-16 | 1.82 | 7.69E-03 | 1.66 |
| **PLK1** | 3.40E-13 | 2.94 | 2.00E-04 | 1.86 | 7.36E-01 | 0.95 | 9.73E-01 | 0.99 | 8.21E-06 | 1.03 | 1.26E-12 | 1.7 | 9.17E-02 | 1.39 |
| **NRAS** | 6.87E-06 | 0.51 | 3.00E-04 | 0.56 | 5.76E-01 | 0.91 | 6.34E-01 | 0.93 | 5.36E-05 | 1.03 | 1.01E-14 | 1.76 | 1.89E-02 | 1.56 |
| **RAD54L** | 6.12E-06 | 2.00 | 5.00E-04 | 1.74 | 9.03E-01 | 1.02 | 6.00E-01 | 0.92 | 3.41E-05 | 1.03 | 7.56E-15 | 1.77 | 1.70E-02 | 1.57 |
| **XRCC2** | 3.24E-06 | 2.04 | 5.00E-04 | 1.73 | 9.81E-01 | 1 | 9.99E-01 | 1 | 1.12E-05 | 1.03 | 1.14E-14 | 1.75 | 2.06E-02 | 1.55 |
| **NFKB2** | 1.31E-06 | 2.08 | 6.00E-04 | 1.73 | 8.21E-01 | 1.04 | 6.96E-01 | 0.94 | 2.47E-05 | 1.03 | 1.40E-14 | 1.76 | 1.21E-02 | 1.61 |
| **E2F1** | 3.78E-06 | 2.04 | 8.00E-04 | 1.73 | 8.99E-01 | 1.02 | 5.51E-01 | 0.91 | 8.59E-06 | 1.03 | 9.14E-14 | 1.73 | 2.01E-02 | 1.55 |
|  | **Kidney (papillary)** | | | | | | | | | | | | | |
|  | **Univariate** | | **Multivariate** | | | | | | | | | | | |
|  | **n=288** | | **n=288** | | **n=288** | | **n=271** | | **n=288** | | **n=259** | | **n=0** | |
| **Gene** | **Expression P-value** | **Expression HR** | **Expression P-value** | **Expression HR** | **Gender P-value** | **Gender HR** | **Race P-value** | **Race HR** | **Age P-value** | **Age HR** | **Stage P-value** | **Stage HR** | **Grade P-value** | **Grade HR** |
| **TGFB1** | 4.57E-04 | 2.78 | 1.00E-04 | 3.8 | 3.33E-01 | 0.71 | 5.71E-01 | 1.13 | 6.19E-01 | 1.01 | 6.16E-09 | 2.63 | No data | No data |
| **PLK1** | 9.54E-11 | 6.25 | 3.00E-04 | 3.88 | 1.11E-01 | 0.56 | 9.29E-01 | 1.02 | 4.62E-01 | 1.01 | 4.14E-05 | 1.99 | No data | No data |
| **DDB2** | 1.11E-08 | 0.21 | 7.00E-04 | 0.29 | 9.80E-01 | 0.99 | 6.26E-01 | 1.11 | 4.19E-01 | 0.99 | 1.96E-06 | 2.1 | No data | No data |
| **RUNX1** | 6.43E-03 | 2.38 | 8.00E-04 | 3.67 | 7.08E-01 | 0.87 | 5.02E-01 | 1.15 | 6.48E-01 | 1.01 | 3.68E-09 | 2.54 | No data | No data |
| **PRC1** | 6.87E-10 | 5.26 | 1.00E-03 | 3.15 | 5.61E-01 | 0.81 | 9.26E-01 | 1.02 | 6.17E-01 | 1.01 | 6.02E-06 | 2.17 | No data | No data |
| **XRCC2** | 1.08E-06 | 4.00 | 1.90E-03 | 2.87 | 3.39E-01 | 0.71 | 7.83E-01 | 1.06 | 5.75E-01 | 1.01 | 1.77E-07 | 2.32 | No data | No data |
| **TRAF1** | 9.70E-02 | 1.69 | 2.50E-03 | 3.28 | 6.70E-02 | 0.49 | 5.71E-01 | 1.13 | 8.68E-01 | 1 | 1.89E-09 | 2.58 | No data | No data |
| **SERPINE1** | 3.41E-03 | 2.78 | 5.90E-03 | 3.07 | 4.05E-01 | 0.74 | 5.23E-01 | 1.15 | 9.67E-01 | 1 | 1.75E-08 | 2.43 | No data | No data |
| **BRIP1** | 2.43E-07 | 4.35 | 8.60E-03 | 2.73 | 7.23E-01 | 0.88 | 3.71E-01 | 1.22 | 5.53E-01 | 1.01 | 3.70E-06 | 2.11 | No data | No data |
| **TOP2A** | 1.27E-09 | 5.26 | 8.60E-03 | 2.64 | 3.47E-01 | 0.71 | 8.40E-01 | 1.04 | 5.63E-01 | 1.01 | 3.86E-05 | 2.03 | No data | No data |
|  | **AML** | | | | | | | | | | | | | |
|  | **Univariate** | | **Multivariate** | | | | | | | | | | | |
|  | **n=151** | | **n=151** | | **n=151** | | **n=149** | | **n=151** | | **n=0** | | **n=0** | |
| **Gene** | **Expression P-value** | **Expression HR** | **Expression P-value** | **Expression HR** | **Gender P-value** | **Gender HR** | **Race P-value** | **Race HR** | **Age P-value** | **Age HR** | **Stage P-value** | **Stage HR** | **Grade P-value** | **Grade HR** |
| **TGFB1** | 5.88E-06 | 2.78 | 9.00E-04 | 2.3 | 2.56E-01 | 0.78 | 8.96E-01 | 1.03 | 1.75E-06 | 1.04 | No data | No data | No data | No data |
| **CDC6** | 8.87E-02 | 1.47 | 4.10E-03 | 1.98 | 5.03E-01 | 0.86 | 6.11E-01 | 1.14 | 1.12E-08 | 1.05 | No data | No data | No data | No data |
| **MAPK1** | 1.56E-03 | 2.00 | 4.20E-03 | 1.91 | 4.58E-01 | 0.85 | 6.17E-01 | 1.14 | 1.76E-07 | 1.04 | No data | No data | No data | No data |
| **RAD51** | 4.73E-02 | 1.56 | 5.80E-03 | 1.87 | 3.99E-01 | 0.83 | 8.28E-01 | 1.06 | 2.39E-08 | 1.05 | No data | No data | No data | No data |
| **EXO1** | 1.65E-02 | 1.75 | 7.20E-03 | 1.95 | 1.87E-01 | 0.74 | 6.73E-01 | 1.11 | 5.12E-08 | 1.05 | No data | No data | No data | No data |
| **MEN1** | 3.75E-03 | 1.92 | 9.10E-03 | 1.85 | 8.30E-01 | 0.95 | 9.98E-01 | 1 | 2.61E-07 | 1.04 | No data | No data | No data | No data |
| **MCM2** | 7.06E-03 | 1.79 | 1.09E-02 | 1.76 | 2.51E-01 | 0.77 | 7.64E-01 | 1.08 | 1.37E-07 | 1.04 | No data | No data | No data | No data |
| **BNIP3** | 1.69E-03 | 0.48 | 1.44E-02 | 0.56 | 4.87E-01 | 0.86 | 7.98E-01 | 1.07 | 4.68E-07 | 1.04 | No data | No data | No data | No data |
|  | **Glioma** | | | | | | | | | | | | | |
|  | **Univariate** | | **Multivariate** | | | | | | | | | | | |
|  | **n=510** | | **n=510** | | **n=510** | | **n=499** | | **n=510** | | **n=0** | | **n=509** | |
| **Gene** | **Expression P-value** | **Expression HR** | **Expression P-value** | **Expression HR** | **Gender P-value** | **Gender HR** | **Race P-value** | **Race HR** | **Age P-value** | **Age HR** | **Stage P-value** | **Stage HR** | **Grade P-value** | **Grade HR** |
| **BNIP3** | 3.00E-11 | 0.31 | 0.00E+00 | 0.32 | 6.31E-01 | 1.09 | 8.38E-01 | 0.96 | 4.04E-13 | 1.06 | No data | No data | 1.68E-06 | 2.69 |
| **BRCA1** | 1.62E-17 | 4.17 | 0.00E+00 | 2.94 | 6.34E-01 | 1.09 | 6.06E-01 | 0.9 | 5.42E-11 | 1.05 | No data | No data | 2.72E-04 | 2.17 |
| **BRIP1** | 6.14E-12 | 3.23 | 0.00E+00 | 2.39 | 8.33E-01 | 1.04 | 8.90E-01 | 0.97 | 4.69E-12 | 1.05 | No data | No data | 1.68E-04 | 2.24 |
| **CDC6** | 1.59E-10 | 3.03 | 0.00E+00 | 2.3 | 7.07E-01 | 1.07 | 7.14E-01 | 0.93 | 1.90E-12 | 1.05 | No data | No data | 9.70E-05 | 2.3 |
| **FBP1** | 3.48E-09 | 2.86 | 0.00E+00 | 3.03 | 9.03E-01 | 1.02 | 8.03E-01 | 0.95 | 8.83E-15 | 1.06 | No data | No data | 2.29E-06 | 2.63 |
| **MCM2** | 4.51E-11 | 3.13 | 0.00E+00 | 2.3 | 5.17E-01 | 1.13 | 7.67E-01 | 1.06 | 4.78E-12 | 1.05 | No data | No data | 7.19E-05 | 2.31 |
| **MYC** | 4.03E-08 | 0.38 | 0.00E+00 | 0.43 | 3.01E-01 | 1.22 | 9.48E-01 | 1.01 | 4.02E-10 | 1.05 | No data | No data | 7.64E-08 | 3 |
| **NRAS** | 5.03E-06 | 3.33 | 0.00E+00 | 3.64 | 4.56E-01 | 1.15 | 9.91E-01 | 1 | 7.99E-15 | 1.06 | No data | No data | 2.10E-06 | 2.63 |
| **RUNX1** | 6.18E-10 | 2.86 | 0.00E+00 | 2.48 | 7.81E-01 | 1.05 | 7.38E-01 | 1.07 | 5.28E-14 | 1.06 | No data | No data | 4.00E-04 | 2.12 |
| **SERPINE1** | 1.54E-12 | 3.33 | 0.00E+00 | 2.51 | 9.26E-01 | 1.02 | 9.93E-01 | 1 | 2.20E-11 | 1.05 | No data | No data | 8.63E-06 | 2.49 |
|  | **Liver** | | | | | | | | | | | | | |
|  | **Univariate** | | **Multivariate** | | | | | | | | | | | |
|  | **n=371** | | **n=371** | | **n=371** | | **n=359** | | **n=370** | | **n=347** | | **n=366** | |
| **Gene** | **Expression P-value** | **Expression HR** | **Expression P-value** | **Expression HR** | **Gender P-value** | **Gender HR** | **Race P-value** | **Race HR** | **Age P-value** | **Age HR** | **Stage P-value** | **Stage HR** | **Grade P-value** | **Grade HR** |
| **CBX3** | 3.23E-08 | 2.78 | 1.00E-04 | 2.41 | 6.46E-01 | 0.91 | 3.23E-01 | 1.21 | 1.51E-01 | 1.01 | 4.77E-05 | 1.59 | 9.10E-01 | 0.98 |
| **CCNB1** | 6.49E-07 | 2.44 | 1.00E-04 | 2.34 | 3.99E-01 | 0.83 | 3.31E-01 | 1.21 | 5.64E-02 | 1.02 | 5.03E-05 | 1.59 | 9.37E-01 | 1.02 |
| **PLK1** | 4.16E-07 | 2.38 | 1.00E-04 | 2.19 | 3.01E-01 | 0.8 | 1.56E-01 | 1.32 | 6.63E-02 | 1.01 | 1.48E-04 | 1.55 | 8.48E-01 | 1.04 |
| **MCM2** | 1.62E-05 | 2.17 | 2.00E-04 | 2.2 | 6.15E-01 | 0.89 | 5.00E-01 | 1.14 | 9.54E-02 | 1.01 | 1.96E-05 | 1.62 | 9.38E-01 | 1.02 |
| **CDC25A** | 7.21E-06 | 2.17 | 3.00E-04 | 2.08 | 2.30E-01 | 0.77 | 2.88E-01 | 1.24 | 7.05E-02 | 1.01 | 6.57E-05 | 1.58 | 8.77E-01 | 1.03 |
| **MEN1** | 3.87E-04 | 1.85 | 3.00E-04 | 2.05 | 4.62E-01 | 0.85 | 4.81E-01 | 1.15 | 7.27E-02 | 1.01 | 8.71E-07 | 1.73 | 7.16E-01 | 1.08 |
| **MSH2** | 1.79E-05 | 2.22 | 5.00E-04 | 2.14 | 6.67E-01 | 0.91 | 3.26E-01 | 1.21 | 6.05E-02 | 1.02 | 1.78E-05 | 1.63 | 9.46E-01 | 1.01 |
| **TOP2A** | 1.49E-05 | 2.13 | 5.00E-04 | 2.07 | 5.05E-01 | 0.86 | 2.76E-01 | 1.24 | 7.36E-02 | 1.01 | 4.49E-05 | 1.59 | 9.14E-01 | 0.98 |
| **CDC25C** | 2.97E-05 | 2.08 | 6.00E-04 | 2.02 | 4.29E-01 | 0.84 | 3.74E-01 | 1.19 | 4.81E-02 | 1.02 | 2.03E-05 | 1.62 | 6.10E-01 | 1.11 |
| **CDK1** | 1.74E-05 | 2.13 | 7.00E-04 | 2.02 | 3.55E-01 | 0.82 | 2.00E-01 | 1.29 | 6.20E-02 | 1.01 | 2.83E-05 | 1.61 | 9.47E-01 | 0.99 |
|  | **Lung (adeno)** | | | | | | | | | | | | | |
|  | **Univariate** | | **Multivariate** | | | | | | | | | | | |
|  | **n=513** | | **n=513** | | **n=513** | | **n=446** | | **n=494** | | **n=505** | | **n=0** | |
| **Gene** | **Expression P-value** | **Expression HR** | **Expression P-value** | **Expression HR** | **Gender P-value** | **Gender HR** | **Race P-value** | **Race HR** | **Age P-value** | **Age HR** | **Stage P-value** | **Stage HR** | **Grade P-value** | **Grade HR** |
| **CDK1** | 7.69E-06 | 1.92 | 0.00E+00 | 2 | 8.31E-01 | 0.97 | 2.87E-01 | 0.86 | 1.82E-01 | 1.01 | 3.27E-08 | 1.53 | No data | No data |
| **PRC1** | 3.45E-06 | 2.00 | 0.00E+00 | 2.04 | 8.96E-01 | 0.98 | 4.51E-01 | 0.9 | 2.50E-01 | 1.01 | 2.42E-07 | 1.49 | No data | No data |
| **PLK1** | 3.80E-06 | 2.22 | 1.00E-04 | 2.16 | 8.35E-01 | 0.97 | 3.08E-01 | 0.87 | 1.01E-01 | 1.01 | 2.12E-07 | 1.51 | No data | No data |
| **CDKN3** | 2.11E-04 | 1.82 | 4.00E-04 | 1.88 | 8.84E-01 | 1.02 | 3.05E-01 | 0.87 | 1.53E-01 | 1.01 | 3.57E-08 | 1.53 | No data | No data |
| **EXO1** | 6.75E-05 | 1.82 | 4.00E-04 | 1.82 | 6.98E-01 | 0.94 | 3.09E-01 | 0.87 | 2.01E-01 | 1.01 | 2.21E-07 | 1.5 | No data | No data |
| **CBX3** | 1.22E-03 | 1.64 | 5.00E-04 | 1.79 | 9.11E-01 | 0.98 | 4.75E-01 | 0.91 | 2.81E-01 | 1.01 | 3.63E-08 | 1.53 | No data | No data |
| **CDC25C** | 6.30E-05 | 1.89 | 6.00E-04 | 1.84 | 9.31E-01 | 0.99 | 2.34E-01 | 0.85 | 1.79E-01 | 1.01 | 1.44E-07 | 1.51 | No data | No data |
| **RAD51** | 5.59E-05 | 1.85 | 7.00E-04 | 1.78 | 7.27E-01 | 0.94 | 3.10E-01 | 0.87 | 3.44E-01 | 1.01 | 1.82E-07 | 1.5 | No data | No data |
| **CHEK1** | 3.10E-04 | 1.69 | 8.00E-04 | 1.73 | 9.71E-01 | 0.99 | 3.03E-01 | 0.87 | 1.48E-01 | 1.01 | 1.07E-07 | 1.52 | No data | No data |
| **FBP1** | 2.09E-05 | 0.53 | 9.00E-04 | 0.58 | 8.62E-01 | 1.03 | 4.36E-01 | 0.9 | 2.08E-01 | 1.01 | 8.91E-08 | 1.52 | No data | No data |
|  | **Lung (squamous)** | | | | | | | | | | | | | |
|  | **Univariate** | | **Multivariate** | | | | | | | | | | | |
|  | **n=501** | | **n=501** | | **n=501** | | **n=388** | | **n=501** | | **n=497** | | **n=0** | |
| **Gene** | **Expression P-value** | **Expression HR** | **Expression P-value** | **Expression HR** | **Gender P-value** | **Gender HR** | **Race P-value** | **Race HR** | **Age P-value** | **Age HR** | **Stage P-value** | **Stage HR** | **Grade P-value** | **Grade HR** |
| **MSH2** | 1.34E-03 | 0.63 | 1.00E-04 | 0.5 | 5.90E-03 | 1.69 | 2.62E-02 | 1.32 | 1.07E-01 | 1.01 | 9.07E-04 | 1.37 | No data | No data |
| **GADD45G** | 1.65E-03 | 1.59 | 7.00E-04 | 1.81 | 3.81E-03 | 1.74 | 2.05E-02 | 1.33 | 3.34E-02 | 1.02 | 2.18E-04 | 1.45 | No data | No data |
| **SERPINE1** | 2.04E-03 | 1.56 | 1.50E-03 | 1.73 | 5.69E-02 | 1.44 | 9.50E-02 | 1.23 | 1.33E-01 | 1.01 | 1.24E-03 | 1.38 | No data | No data |
| **CDKN3** | 2.21E-02 | 0.72 | 2.20E-03 | 0.6 | 7.72E-03 | 1.65 | 2.02E-02 | 1.33 | 8.92E-02 | 1.02 | 3.26E-04 | 1.43 | No data | No data |
| **EREG** | 1.24E-03 | 1.56 | 2.30E-03 | 1.63 | 1.53E-02 | 1.58 | 1.06E-01 | 1.23 | 6.36E-02 | 1.02 | 2.07E-03 | 1.35 | No data | No data |
| **XRCC2** | 2.38E-02 | 0.71 | 4.10E-03 | 0.61 | 9.27E-03 | 1.64 | 1.41E-02 | 1.36 | 1.04E-01 | 1.01 | 1.71E-03 | 1.36 | No data | No data |
| **TOP2A** | 5.18E-03 | 0.68 | 5.10E-03 | 0.63 | 1.42E-02 | 1.58 | 5.42E-02 | 1.27 | 1.55E-01 | 1.01 | 1.04E-03 | 1.37 | No data | No data |
| **MAPK1** | 3.07E-02 | 0.74 | 6.50E-03 | 0.65 | 1.37E-02 | 1.59 | 2.99E-02 | 1.31 | 5.09E-02 | 1.02 | 1.42E-03 | 1.36 | No data | No data |
| **PLK1** | 1.44E-02 | 0.71 | 7.30E-03 | 0.64 | 1.21E-02 | 1.61 | 2.99E-02 | 1.31 | 1.64E-01 | 1.01 | 1.43E-03 | 1.36 | No data | No data |
| **DDB2** | 4.83E-03 | 0.68 | 1.12E-02 | 0.67 | 1.72E-02 | 1.57 | 2.83E-02 | 1.31 | 5.15E-02 | 1.02 | 1.19E-03 | 1.38 | No data | No data |
|  | **Ovarium** | | | | | | | | | | | | | |
|  | **Univariate** | | **Multivariate** | | | | | | | | | | | |
|  | **n=374** | | **n=374** | | **n=374** | | **n=360** | | **n=374** | | **n=0** | | **n=364** | |
| **Gene** | **Expression P-value** | **Expression HR** | **Expression P-value** | **Expression HR** | **Gender P-value** | **Gender HR** | **Race P-value** | **Race HR** | **Age P-value** | **Age HR** | **Stage P-value** | **Stage HR** | **Grade P-value** | **Grade HR** |
| **MYC** | 5.83E-04 | 1.69 | 1.00E-04 | 1.9 | - | - | 2.60E-01 | 1.16 | 8.90E-05 | 1.03 | No data | No data | 5.26E-01 | 1.14 |
| **CBX3** | 1.48E-03 | 0.65 | 4.00E-04 | 0.61 | - | - | 2.60E-02 | 1.33 | 8.27E-04 | 1.02 | No data | No data | 6.34E-01 | 1.11 |
| **RUNX1** | 4.30E-04 | 1.59 | 1.20E-03 | 1.56 | - | - | 1.16E-01 | 1.22 | 1.38E-03 | 1.02 | No data | No data | 6.21E-01 | 1.11 |
| **DDB2** | 4.06E-03 | 0.64 | 9.70E-03 | 0.66 | - | - | 6.74E-02 | 1.26 | 4.94E-03 | 1.02 | No data | No data | 2.96E-01 | 1.25 |
| **CXCL9** | 8.25E-04 | 0.63 | 9.90E-03 | 0.69 | - | - | 1.56E-01 | 1.2 | 2.48E-03 | 1.02 | No data | No data | 4.00E-01 | 1.19 |
| **NRAS** | 3.87E-03 | 0.68 | 1.32E-02 | 0.71 | - | - | 5.40E-02 | 1.28 | 1.33E-03 | 1.02 | No data | No data | 5.72E-01 | 1.13 |
| **GADD45G** | 5.10E-03 | 0.69 | 1.36E-02 | 0.71 | - | - | 9.10E-02 | 1.24 | 2.90E-03 | 1.02 | No data | No data | 3.45E-01 | 1.22 |
| **FANCD2** | 1.60E-02 | 1.45 | 1.65E-02 | 1.47 | - | - | 1.69E-01 | 1.19 | 8.54E-04 | 1.02 | No data | No data | 4.98E-01 | 1.15 |
| **XRCC2** | 9.51E-03 | 0.69 | 3.40E-02 | 0.73 | - | - | 4.58E-02 | 1.29 | 2.09E-03 | 1.02 | No data | No data | 3.56E-01 | 1.22 |
| **CCNB1** | 1.87E-02 | 0.73 | 3.82E-02 | 0.75 | - | - | 5.10E-02 | 1.28 | 2.13E-03 | 1.02 | No data | No data | 4.17E-01 | 1.19 |
|  | **Pancreas** | | | | | | | | | | | | | |
|  | **Univariate** | | **Multivariate** | | | | | | | | | | | |
|  | **n=177** | | **n=177** | | **n=177** | | **n=173** | | **n=177** | | **n=174** | | **n=175** | |
| **Gene** | **Expression P-value** | **Expression HR** | **Expression P-value** | **Expression HR** | **Gender P-value** | **Gender HR** | **Race P-value** | **Race HR** | **Age P-value** | **Age HR** | **Stage P-value** | **Stage HR** | **Grade P-value** | **Grade HR** |
| **CDK1** | 4.10E-06 | 2.63 | 0.00E+00 | 2.53 | 4.77E-01 | 0.86 | 7.58E-01 | 1.08 | 5.36E-02 | 1.02 | 1.71E-01 | 1.33 | 1.20E-01 | 1.43 |
| **TRAF1** | 2.21E-05 | 0.42 | 0.00E+00 | 0.35 | 1.19E-01 | 0.71 | 6.23E-01 | 1.12 | 1.00E-01 | 1.02 | 1.23E-01 | 1.37 | 4.31E-02 | 1.6 |
| **E2F1** | 4.18E-04 | 2.04 | 3.00E-04 | 2.23 | 5.31E-01 | 0.87 | 9.98E-01 | 1 | 1.30E-01 | 1.02 | 9.40E-02 | 1.44 | 8.64E-02 | 1.48 |
| **TOP2A** | 4.88E-05 | 2.50 | 4.00E-04 | 2.36 | 7.11E-01 | 0.92 | 9.88E-01 | 1 | 7.60E-02 | 1.02 | 4.69E-01 | 1.17 | 2.07E-01 | 1.34 |
| **GADD45G** | 3.76E-05 | 0.33 | 7.00E-04 | 0.38 | 5.87E-01 | 0.89 | 6.42E-01 | 0.89 | 4.08E-02 | 1.02 | 6.87E-01 | 1.09 | 1.49E-01 | 1.39 |
| **CBX3** | 2.18E-04 | 2.33 | 1.00E-03 | 2.2 | 4.64E-01 | 0.85 | 7.64E-01 | 1.08 | 6.71E-02 | 1.02 | 4.59E-01 | 1.17 | 1.23E-01 | 1.42 |
| **CHEK1** | 6.07E-06 | 2.63 | 1.10E-03 | 2.25 | 8.37E-01 | 1.05 | 7.41E-01 | 1.08 | 7.39E-02 | 1.02 | 2.70E-01 | 1.26 | 3.32E-01 | 1.25 |
| **PRC1** | 5.25E-05 | 2.33 | 1.40E-03 | 2.03 | 6.80E-01 | 0.91 | 5.68E-01 | 1.15 | 5.69E-02 | 1.02 | 2.43E-01 | 1.28 | 2.02E-01 | 1.34 |
| **SERPINE1** | 2.87E-03 | 1.89 | 1.50E-03 | 2.02 | 5.22E-01 | 0.87 | 9.73E-01 | 0.99 | 2.22E-02 | 1.03 | 1.61E-01 | 1.34 | 1.61E-01 | 1.38 |
| **EREG** | 2.44E-04 | 2.22 | 1.60E-03 | 2.09 | 4.47E-01 | 0.85 | 5.73E-01 | 1.14 | 1.74E-02 | 1.03 | 3.08E-01 | 1.22 | 1.98E-01 | 1.34 |
|  | **Paraganglioma** | | | | | | | | | | | | | |
|  | **Univariate** | | **Multivariate** | | | | | | | | | | | |
|  | **n=178** | | **n=178** | | **n=178** | | **n=173** | | **n=178** | | **n=0** | | **n=0** | |
| **Gene** | **Expression P-value** | **Expression HR** | **Expression P-value** | **Expression HR** | **Gender P-value** | **Gender HR** | **Race P-value** | **Race HR** | **Age P-value** | **Age HR** | **Stage P-value** | **Stage HR** | **Grade P-value** | **Grade HR** |
| **FANCD2** | 1.18E-03 | 14.29 | 6.90E-03 | 20.66 | 1.39E-01 | 4.3 | 9.99E-01 | 0 | 6.49E-02 | 1.05 | No data | No data | No data | No data |
| **CDK1** | 3.79E-03 | 12.50 | 8.20E-03 | 19.93 | 1.48E-01 | 3.98 | 9.99E-01 | 0 | 8.78E-02 | 1.05 | No data | No data | No data | No data |
| **MEN1** | 1.74E-03 | 14.29 | 1.52E-02 | 16.41 | 1.74E-01 | 4.51 | 9.99E-01 | 0 | 1.93E-01 | 1.05 | No data | No data | No data | No data |
| **CCNB1** | 1.25E-02 | 10.00 | 2.42E-02 | 12.65 | 1.46E-01 | 4.04 | 9.99E-01 | 0 | 1.26E-01 | 1.05 | No data | No data | No data | No data |
| **PRC1** | 1.92E-02 | 9.09 | 2.89E-02 | 12.26 | 1.99E-01 | 3.7 | 9.99E-01 | 0 | 9.33E-02 | 1.05 | No data | No data | No data | No data |
| **CBX3** | 2.55E-02 | 5.56 | 3.01E-02 | 6.93 | 1.02E-01 | 6.08 | 9.99E-01 | 0 | 2.91E-01 | 1.03 | No data | No data | No data | No data |
| **DDB2** | 4.85E-02 | 5.00 | 3.91E-02 | 6.45 | 1.18E-01 | 4.71 | 9.99E-01 | 0 | 1.96E-01 | 1.04 | No data | No data | No data | No data |
| **SERPINE1** | 1.98E-02 | 0.12 | 4.53E-02 | 0.11 | 9.61E-02 | 6.06 | 9.99E-01 | 0 | 4.45E-01 | 1.02 | No data | No data | No data | No data |
| **E2F1** | 4.74E-02 | 5.00 | 4.87E-02 | 5.9 | 1.30E-01 | 4.3 | 9.99E-01 | 0 | 1.46E-01 | 1.04 | No data | No data | No data | No data |
|  | **Prostate** | | | | | | | | | | | | | |
|  | **Univariate** | | **Multivariate** | | | | | | | | | | | |
|  | **n=495** | | **n=495** | | **n=0** | | **n=156** | | **n=495** | | **n=0** | | **n=0** | |
| **Gene** | **Expression P-value** | **Expression HR** | **Expression P-value** | **Expression HR** | **Gender P-value** | **Gender HR** | **Race P-value** | **Race HR** | **Age P-value** | **Age HR** | **Stage P-value** | **Stage HR** | **Grade P-value** | **Grade HR** |
| **NEIL1** | 4.08E-03 | 7.69 | 0.00E+00 | >100 | No data | No data | 0.00E+00 | 1 | 0.00E+00 | 1 | No data | No data | No data | No data |
| **BNIP3** | 2.92E-02 | 8.33 | 9.99E-01 | >100 | No data | No data | 1.00E+00 | 0 | 5.58E-01 | 0.92 | No data | No data | No data | No data |
| **FANCD2** | 1.66E-01 | 3.85 | 9.99E-01 | >100 | No data | No data | 1.00E+00 | 0 | 7.01E-01 | 0.95 | No data | No data | No data | No data |
| **FBP1** | 1.50E-01 | 0.33 | 9.99E-01 | >100 | No data | No data | 1.00E+00 | 0 | 4.80E-01 | 0.87 | No data | No data | No data | No data |
| **MCM2** | 1.75E-02 | 5.26 | 9.99E-01 | >100 | No data | No data | 1.00E+00 | 0 | 6.42E-01 | 0.94 | No data | No data | No data | No data |
| **MCM4** | 8.29E-02 | 5.26 | 9.99E-01 | >100 | No data | No data | 1.00E+00 | 0 | 4.74E-01 | 0.9 | No data | No data | No data | No data |
| **RUNX1** | 3.64E-02 | 0.22 | 9.99E-01 | 0 | No data | No data | 1.00E+00 | 0 | 5.84E-01 | 0.92 | No data | No data | No data | No data |
| **TGFB1** | 1.96E-01 | 0.37 | 9.99E-01 | 0 | No data | No data | 1.00E+00 | 0 | 6.72E-01 | 0.94 | No data | No data | No data | No data |
| **TOPBP1** | 6.97E-03 | 5.00 | 9.99E-01 | >100 | No data | No data | 1.00E+00 | 0 | 5.50E-01 | 0.9 | No data | No data | No data | No data |
| **BRCA1** | 1.34E-01 | 4.35 | 1.00E+00 | >100 | No data | No data | 1.00E+00 | 0 | 6.49E-01 | 0.94 | No data | No data | No data | No data |
|  | **Rectum** | | | | | | | | | | | | | |
|  | **Univariate** | | **Multivariate** | | | | | | | | | | | |
|  | **n=165** | | **n=165** | | **n=165** | | **n=87** | | **n=165** | | **n=156** | | **n=0** | |
| **Gene** | **Expression P-value** | **Expression HR** | **Expression P-value** | **Expression HR** | **Gender P-value** | **Gender HR** | **Race P-value** | **Race HR** | **Age P-value** | **Age HR** | **Stage P-value** | **Stage HR** | **Grade P-value** | **Grade HR** |
| **NRAS** | 8.97E-03 | 0.36 | 3.95E-02 | 0.25 | 7.61E-01 | 1.23 | 2.07E-01 | 2.33 | 1.84E-02 | 1.12 | 6.51E-01 | 1.2 | No data | No data |
| **EREG** | 7.02E-03 | 0.32 | 4.60E-02 | 0.2 | 5.03E-01 | 0.68 | 1.30E-01 | 3.04 | 7.62E-03 | 1.15 | 4.81E-01 | 1.3 | No data | No data |
| **TGFB1** | 3.60E-01 | 1.47 | 4.64E-02 | 3.86 | 6.21E-01 | 0.74 | 4.44E-01 | 1.66 | 7.44E-03 | 1.15 | 6.15E-01 | 1.21 | No data | No data |
| **FBP1** | 8.96E-02 | 1.96 | 7.79E-02 | 3.11 | 9.43E-01 | 1.05 | 2.02E-01 | 2.32 | 1.16E-02 | 1.13 | 5.59E-01 | 1.25 | No data | No data |
| **RAD54L** | 2.70E-02 | 0.40 | 9.29E-02 | 0.31 | 4.64E-01 | 0.65 | 2.07E-01 | 2.37 | 6.57E-03 | 1.15 | 8.95E-01 | 1.05 | No data | No data |
| **NFKB2** | 2.72E-02 | 2.56 | 1.07E-01 | 3.29 | 6.08E-01 | 0.73 | 5.67E-01 | 1.47 | 2.09E-02 | 1.11 | 7.83E-01 | 0.9 | No data | No data |
| **GADD45G** | 3.83E-03 | 6.25 | 1.10E-01 | 5.57 | 4.99E-01 | 0.67 | 3.03E-01 | 1.96 | 2.65E-02 | 1.1 | 6.40E-01 | 1.18 | No data | No data |
| **E2F1** | 2.63E-01 | 0.63 | 1.70E-01 | 0.39 | 5.57E-01 | 0.71 | 2.05E-01 | 2.42 | 7.29E-03 | 1.15 | 7.52E-01 | 1.12 | No data | No data |
| **CDKN3** | 1.96E-01 | 1.75 | 1.86E-01 | 2.42 | 7.84E-01 | 0.85 | 3.66E-01 | 1.77 | 8.63E-03 | 1.14 | 6.46E-01 | 1.19 | No data | No data |
| **BRCA1** | 3.40E-04 | 0.25 | 2.03E-01 | 0.43 | 8.13E-01 | 0.87 | 3.13E-01 | 1.94 | 3.60E-02 | 1.11 | 7.89E-01 | 1.11 | No data | No data |
|  | **Sarcoma** | | | | | | | | | | | | | |
|  | **Univariate** | | **Multivariate** | | | | | | | | | | | |
|  | **n=259** | | **n=259** | | **n=259** | | **n=250** | | **n=259** | | **n=0** | | **n=0** | |
| **Gene** | **Expression P-value** | **Expression HR** | **Expression P-value** | **Expression HR** | **Gender P-value** | **Gender HR** | **Race P-value** | **Race HR** | **Age P-value** | **Age HR** | **Stage P-value** | **Stage HR** | **Grade P-value** | **Grade HR** |
| **BNIP3** | 2.19E-05 | 2.38 | 1.00E-04 | 2.32 | 3.96E-01 | 0.83 | 6.30E-01 | 1.1 | 1.01E-02 | 1.02 | No data | No data | No data | No data |
| **NFKB2** | 5.68E-05 | 0.42 | 2.00E-04 | 0.42 | 4.39E-01 | 0.85 | 2.22E-01 | 1.28 | 1.66E-02 | 1.02 | No data | No data | No data | No data |
| **MSH2** | 5.87E-04 | 2.38 | 3.00E-04 | 2.68 | 7.02E-01 | 1.09 | 1.48E-01 | 1.35 | 1.80E-03 | 1.03 | No data | No data | No data | No data |
| **CDC25A** | 1.01E-04 | 2.38 | 4.00E-04 | 2.36 | 9.01E-01 | 0.97 | 5.68E-01 | 1.12 | 1.91E-02 | 1.02 | No data | No data | No data | No data |
| **RAD54L** | 3.78E-04 | 2.04 | 5.00E-04 | 2.13 | 6.89E-01 | 0.92 | 6.60E-01 | 1.09 | 6.39E-03 | 1.02 | No data | No data | No data | No data |
| **CXCL9** | 4.49E-03 | 0.52 | 6.00E-04 | 0.43 | 3.84E-01 | 0.83 | 4.24E-01 | 1.17 | 1.61E-03 | 1.02 | No data | No data | No data | No data |
| **TOP2A** | 6.14E-04 | 2.13 | 1.10E-03 | 2.19 | 7.72E-01 | 1.07 | 2.54E-01 | 1.26 | 1.08E-02 | 1.02 | No data | No data | No data | No data |
| **CHEK1** | 1.22E-03 | 1.92 | 1.20E-03 | 2.02 | 9.65E-01 | 0.99 | 2.79E-01 | 1.25 | 4.40E-03 | 1.02 | No data | No data | No data | No data |
| **FBP1** | 4.74E-03 | 0.49 | 1.40E-03 | 0.42 | 8.10E-01 | 0.95 | 1.70E-01 | 1.32 | 3.62E-03 | 1.02 | No data | No data | No data | No data |
| **MCM4** | 6.94E-04 | 2.04 | 1.50E-03 | 2.03 | 9.70E-01 | 0.99 | 3.45E-01 | 1.21 | 9.12E-03 | 1.02 | No data | No data | No data | No data |
|  | **Melanoma** | | | | | | | | | | | | | |
|  | **Univariate** | | **Multivariate** | | | | | | | | | | | |
|  | **n=468** | | **n=468** | | **n=468** | | **n=458** | | **n=460** | | **n=416** | | **n=0** | |
| **Gene** | **Expression P-value** | **Expression HR** | **Expression P-value** | **Expression HR** | **Gender P-value** | **Gender HR** | **Race P-value** | **Race HR** | **Age P-value** | **Age HR** | **Stage P-value** | **Stage HR** | **Grade P-value** | **Grade HR** |
| **CXCL9** | 2.74E-08 | 0.40 | 0.00E+00 | 0.41 | 9.96E-01 | 1 | 4.83E-03 | 3.07 | 8.96E-04 | 1.02 | 2.66E-04 | 1.41 | No data | No data |
| **TRAF1** | 1.78E-06 | 0.52 | 0.00E+00 | 0.47 | 8.07E-01 | 0.96 | 1.62E-03 | 3.51 | 4.30E-05 | 1.02 | 4.11E-05 | 1.46 | No data | No data |
| **CDC25A** | 1.09E-05 | 1.82 | 1.00E-04 | 1.84 | 9.77E-01 | 1 | 2.32E-04 | 4.33 | 2.36E-04 | 1.02 | 4.90E-04 | 1.37 | No data | No data |
| **FBP1** | 1.43E-05 | 0.55 | 1.00E-04 | 0.55 | 9.09E-01 | 0.98 | 3.19E-03 | 3.24 | 1.84E-04 | 1.02 | 1.25E-04 | 1.42 | No data | No data |
| **PRC1** | 9.01E-05 | 1.72 | 2.00E-04 | 1.78 | 9.66E-01 | 0.99 | 1.62E-04 | 4.54 | 4.08E-05 | 1.02 | 2.52E-04 | 1.38 | No data | No data |
| **FANCD2** | 8.94E-03 | 1.45 | 3.00E-04 | 1.75 | 8.37E-01 | 0.97 | 5.77E-04 | 3.92 | 5.16E-05 | 1.02 | 1.05E-04 | 1.42 | No data | No data |
| **NFKB2** | 2.53E-04 | 0.59 | 3.00E-04 | 0.57 | 9.84E-01 | 1 | 7.11E-04 | 3.83 | 9.95E-05 | 1.02 | 4.36E-04 | 1.38 | No data | No data |
| **CDC25C** | 7.90E-03 | 1.47 | 4.00E-04 | 1.74 | 9.03E-01 | 1.02 | 5.65E-04 | 3.93 | 1.07E-04 | 1.02 | 9.59E-05 | 1.42 | No data | No data |
| **PLK1** | 1.65E-04 | 1.67 | 4.00E-04 | 1.7 | 9.64E-01 | 1.01 | 1.84E-03 | 3.45 | 3.02E-05 | 1.02 | 2.60E-04 | 1.39 | No data | No data |
| **GADD45G** | 7.69E-03 | 0.69 | 6.00E-04 | 0.6 | 7.17E-01 | 0.94 | 8.79E-04 | 3.76 | 1.15E-05 | 1.02 | 1.62E-04 | 1.42 | No data | No data |
|  | **Stomach** | | | | | | | | | | | | | |
|  | **Univariate** | | **Multivariate** | | | | | | | | | | | |
|  | **n=375** | | **n=375** | | **n=375** | | **n=323** | | **n=371** | | **n=352** | | **n=366** | |
| **Gene** | **Expression P-value** | **Expression HR** | **Expression P-value** | **Expression HR** | **Gender P-value** | **Gender HR** | **Race P-value** | **Race HR** | **Age P-value** | **Age HR** | **Stage P-value** | **Stage HR** | **Grade P-value** | **Grade HR** |
| **MEN1** | 2.46E-05 | 0.48 | 1.00E-04 | 0.45 | 2.24E-01 | 1.28 | 6.38E-01 | 1.09 | 8.62E-04 | 1.03 | 1.43E-03 | 1.55 | 1.30E-01 | 1.37 |
| **BNIP3** | 1.98E-03 | 1.96 | 8.00E-04 | 2.49 | 1.90E-01 | 1.31 | 8.90E-01 | 1.02 | 3.16E-03 | 1.03 | 1.42E-03 | 1.54 | 1.97E-01 | 1.31 |
| **SERPINE1** | 5.35E-06 | 2.13 | 8.00E-04 | 1.95 | 4.48E-01 | 1.17 | 4.66E-01 | 1.14 | 7.14E-03 | 1.03 | 3.19E-03 | 1.5 | 1.22E-01 | 1.38 |
| **DDB2** | 1.28E-02 | 0.62 | 3.10E-03 | 0.5 | 2.09E-01 | 1.3 | 9.25E-01 | 1.02 | 3.37E-03 | 1.03 | 8.53E-04 | 1.58 | 3.48E-02 | 1.56 |
| **TRAF1** | 7.87E-03 | 0.62 | 3.10E-03 | 0.51 | 2.86E-01 | 1.25 | 9.52E-01 | 0.99 | 1.65E-03 | 1.03 | 4.54E-04 | 1.62 | 3.62E-02 | 1.57 |
| **RAD51** | 2.52E-02 | 0.68 | 5.90E-03 | 0.57 | 3.39E-01 | 1.22 | 7.38E-01 | 1.06 | 1.80E-03 | 1.03 | 1.33E-03 | 1.55 | 1.31E-01 | 1.37 |
| **BRIP1** | 9.50E-03 | 0.57 | 7.80E-03 | 0.5 | 1.83E-01 | 1.32 | 5.77E-01 | 1.1 | 1.86E-03 | 1.03 | 1.78E-03 | 1.53 | 1.16E-01 | 1.39 |
| **BRCA1** | 1.33E-03 | 0.59 | 8.70E-03 | 0.59 | 2.10E-01 | 1.29 | 5.37E-01 | 1.12 | 1.68E-03 | 1.03 | 1.44E-03 | 1.55 | 1.74E-01 | 1.33 |
| **MCM2** | 1.02E-02 | 0.65 | 1.03E-02 | 0.6 | 2.21E-01 | 1.29 | 7.37E-01 | 1.06 | 2.45E-03 | 1.03 | 8.96E-04 | 1.58 | 2.00E-01 | 1.31 |
| **TGFB1** | 1.29E-03 | 1.69 | 1.05E-02 | 1.66 | 2.51E-01 | 1.27 | 7.73E-01 | 1.05 | 4.38E-03 | 1.03 | 7.13E-04 | 1.59 | 2.53E-01 | 1.27 |
|  | **Testis** | | | | | | | | | | | | | |
|  | **Univariate** | | **Multivariate** | | | | | | | | | | | |
|  | **n=134** | | **n=134** | | **n=134** | | **n=129** | | **n=134** | | **n=81** | | **n=0** | |
| **Gene** | **Expression P-value** | **Expression HR** | **Expression P-value** | **Expression HR** | **Gender P-value** | **Gender HR** | **Race P-value** | **Race HR** | **Age P-value** | **Age HR** | **Stage P-value** | **Stage HR** | **Grade P-value** | **Grade HR** |
| **CXCL9** | 1.08E-01 | 5.56 | 1.00E+00 | >100 | - | - | 1.00E+00 | >100 | 9.99E-01 | 0.01 | 1.00E+00 | 0.39 | No data | No data |
| **DDB2** | 4.25E-02 | 7.69 | 1.00E+00 | 0 | - | - | 1.00E+00 | >100 | 9.99E-01 | 0.01 | 1.00E+00 | 0.39 | No data | No data |
| **MEN1** | 1.34E-01 | 1.00 | 1.00E+00 | 0 | - | - | 1.00E+00 | >100 | 9.99E-01 | 0.01 | 1.00E+00 | 0.34 | No data | No data |
| **RUNX1** | 1.84E-01 | 3.45 | 1.00E+00 | 0 | - | - | 1.00E+00 | >100 | 9.99E-01 | 0.01 | 1.00E+00 | 0.57 | No data | No data |
| **CBX3** | 4.69E-02 | 0.13 | 1.00E+00 | 0 | - | - | 1.00E+00 | >100 | 1.00E+00 | 0.5 | 1.00E+00 | >100 | No data | No data |
| **CDC6** | 1.35E-01 | 4.76 | 1.00E+00 | 0 | - | - | 1.00E+00 | >100 | 9.99E-01 | 0.01 | 1.00E+00 | >100 | No data | No data |
| **CDKN3** | 9.46E-01 | 1.09 | 1.00E+00 | >100 | - | - | 1.00E+00 | >100 | 9.99E-01 | 0.01 | 1.00E+00 | 0.27 | No data | No data |
| **FBP1** | 3.50E-02 | 1.00 | 1.00E+00 | >100 | - | - | 1.00E+00 | >100 | 9.99E-01 | 0.16 | 1.00E+00 | >100 | No data | No data |
| **MAPK1** | 1.94E-01 | 4.17 | 1.00E+00 | 0 | - | - | 1.00E+00 | >100 | 9.99E-01 | 0.01 | 1.00E+00 | >100 | No data | No data |
| **MCM4** | 2.78E-01 | 3.23 | 1.00E+00 | >100 | - | - | 1.00E+00 | >100 | 9.99E-01 | 0.16 | 1.00E+00 | >100 | No data | No data |
|  | **Thyroid** | | | | | | | | | | | | | |
|  | **Univariate** | | **Multivariate** | | | | | | | | | | | |
|  | **n=502** | | **n=502** | | **n=502** | | **n=410** | | **n=502** | | **n=500** | | **n=0** | |
| **Gene** | **Expression P-value** | **Expression HR** | **Expression P-value** | **Expression HR** | **Gender P-value** | **Gender HR** | **Race P-value** | **Race HR** | **Age P-value** | **Age HR** | **Stage P-value** | **Stage HR** | **Grade P-value** | **Grade HR** |
| **BNIP3** | 5.00E-02 | 2.56 | 1.11E-02 | 4.27 | 7.31E-01 | 0.82 | 6.18E-01 | 1.23 | 6.80E-07 | 1.13 | 2.75E-01 | 1.39 | No data | No data |
| **CDC6** | 1.82E-02 | 0.32 | 1.65E-02 | 0.27 | 8.48E-01 | 1.11 | 5.05E-01 | 1.32 | 2.70E-07 | 1.13 | 5.27E-01 | 1.19 | No data | No data |
| **RAD51** | 2.65E-01 | 0.56 | 3.32E-02 | 0.29 | 8.47E-01 | 1.11 | 6.52E-01 | 1.2 | 2.34E-07 | 1.13 | 3.47E-01 | 1.3 | No data | No data |
| **E2F1** | 1.59E-04 | 0.15 | 4.20E-02 | 0.29 | 6.67E-01 | 0.79 | 4.97E-01 | 1.33 | 2.89E-06 | 1.1 | 4.62E-01 | 1.25 | No data | No data |
| **BRIP1** | 4.90E-02 | 0.38 | 4.71E-02 | 0.36 | 9.89E-01 | 0.99 | 5.82E-01 | 1.26 | 6.90E-07 | 1.12 | 3.49E-01 | 1.31 | No data | No data |
|  | **Thymoma** | | | | | | | | | | | | | |
|  | **Univariate** | | **Multivariate** | | | | | | | | | | | |
|  | **n=119** | | **n=119** | | **n=119** | | **n=117** | | **n=118** | | **n=0** | | **n=0** | |
| **Gene** | **Expression P-value** | **Expression HR** | **Expression P-value** | **Expression HR** | **Gender P-value** | **Gender HR** | **Race P-value** | **Race HR** | **Age P-value** | **Age HR** | **Stage P-value** | **Stage HR** | **Grade P-value** | **Grade HR** |
| **MSH2** | 8.50E-04 | 0.12 | 5.20E-03 | 0.07 | 8.76E-01 | 0.89 | 7.53E-01 | 0.72 | 3.73E-02 | 1.08 | No data | No data | No data | No data |
| **TRAF1** | 1.36E-02 | 5.88 | 8.70E-03 | 11.1 | 6.48E-01 | 1.39 | 6.77E-01 | 0.66 | 9.89E-03 | 1.08 | No data | No data | No data | No data |
| **BRCA1** | 3.36E-04 | 0.06 | 1.05E-02 | 0.06 | 6.23E-01 | 0.67 | 3.03E-01 | 0.3 | 4.49E-01 | 1.02 | No data | No data | No data | No data |
| **FANCD2** | 5.98E-04 | 0.10 | 1.15E-02 | 0.11 | 8.28E-01 | 0.85 | 6.51E-01 | 0.63 | 2.32E-01 | 1.04 | No data | No data | No data | No data |
| **BRIP1** | 1.09E-04 | 0.04 | 1.23E-02 | 0.05 | 8.77E-01 | 1.12 | 4.39E-01 | 0.43 | 5.00E-01 | 1.02 | No data | No data | No data | No data |
| **CDK1** | 5.36E-04 | 0.10 | 1.34E-02 | 0.07 | 8.72E-01 | 0.89 | 2.12E-01 | 0.21 | 8.38E-01 | 1.01 | No data | No data | No data | No data |
| **RAD51** | 6.93E-04 | 0.10 | 1.41E-02 | 0.06 | 9.07E-01 | 1.09 | 1.69E-01 | 0.17 | 8.16E-01 | 1.01 | No data | No data | No data | No data |
| **CCNB1** | 7.91E-04 | 0.10 | 1.47E-02 | 0.07 | 8.70E-01 | 0.88 | 2.10E-01 | 0.21 | 7.48E-01 | 1.01 | No data | No data | No data | No data |
| **CXCL9** | 1.76E-03 | 7.14 | 1.70E-02 | 7.07 | 6.84E-01 | 1.36 | 5.06E-01 | 0.51 | 1.46E-01 | 1.04 | No data | No data | No data | No data |
| **TOPBP1** | 2.54E-03 | 0.12 | 1.72E-02 | 0.13 | 9.47E-01 | 0.95 | 6.04E-01 | 0.59 | 1.21E-01 | 1.05 | No data | No data | No data | No data |
|  | **Uterine** | | | | | | | | | | | | | |
|  | **Univariate** | | **Multivariate** | | | | | | | | | | | |
|  | **n=543** | | **n=543** | | **n=543** | | **n=498** | | **n=540** | | **n=0** | | **n=543** | |
| **Gene** | **Expression P-value** | **Expression HR** | **Expression P-value** | **Expression HR** | **Gender P-value** | **Gender HR** | **Race P-value** | **Race HR** | **Age P-value** | **Age HR** | **Stage P-value** | **Stage HR** | **Grade P-value** | **Grade HR** |
| **CXCL9** | 5.17E-04 | 0.48 | 9.00E-04 | 0.48 | - | - | 9.06E-01 | 1.02 | 4.70E-03 | 1.03 | No data | No data | 1.95E-05 | 3.32 |
| **MCM4** | 5.62E-05 | 2.56 | 4.70E-03 | 2.12 | - | - | 9.56E-01 | 0.99 | 2.16E-03 | 1.03 | No data | No data | 1.55E-03 | 2.49 |
| **MCM2** | 1.27E-03 | 1.96 | 1.65E-02 | 1.72 | - | - | 8.52E-01 | 0.98 | 1.33E-03 | 1.03 | No data | No data | 7.98E-04 | 2.64 |
| **SERPINE1** | 3.26E-02 | 1.79 | 1.72E-02 | 2.02 | - | - | 5.87E-01 | 1.08 | 1.71E-03 | 1.03 | No data | No data | 1.22E-04 | 2.94 |
| **TRAF1** | 5.59E-03 | 0.43 | 2.64E-02 | 0.49 | - | - | 8.79E-01 | 1.02 | 3.79E-03 | 1.03 | No data | No data | 1.24E-04 | 2.94 |
| **NFKB2** | 2.15E-02 | 0.62 | 3.19E-02 | 0.63 | - | - | 9.19E-01 | 0.99 | 3.36E-03 | 1.03 | No data | No data | 4.68E-05 | 3.12 |
| **E2F1** | 2.11E-05 | 2.78 | 3.48E-02 | 1.85 | - | - | 8.96E-01 | 0.98 | 5.66E-03 | 1.03 | No data | No data | 5.41E-03 | 2.35 |
| **MSH2** | 5.52E-04 | 2.04 | 4.01E-02 | 1.6 | - | - | 9.69E-01 | 1.01 | 3.50E-03 | 1.03 | No data | No data | 1.10E-03 | 2.61 |
| **BRIP1** | 1.08E-03 | 2.00 | 4.44E-02 | 1.59 | - | - | 7.86E-01 | 1.04 | 2.39E-03 | 1.03 | No data | No data | 3.82E-04 | 2.77 |
| **EREG** | 6.62E-02 | 1.47 | 4.80E-02 | 1.53 | - | - | 8.73E-01 | 1.02 | 1.30E-03 | 1.03 | No data | No data | 6.13E-05 | 3.07 |
